# GDF3 is an endogenous antagonist of the ActE-ALK7/ACVR2 pathway in adipocytes

**DOI:** 10.64898/2026.09.18.752693

**Authors:** Nagasuryaprasad Kotikalapudi, Deepti Ramachandran, Evelyn Y. Zhou, James A. Howard, Daniel E. Vieira, Wei-An Chen, Sarah Hsu, Kyung Cheul Shin, Shirish Naz, Z. Gordon Jiang, Josep M. Mercader, Thomas B. Thompson, Stephen E. Flaherty, Paul M. Titchenell, Henning F. Kramer, Alexander S. Banks

## Abstract

Metabolic dysfunction-associated steatotic liver disease (MASLD) and steatohepatitis (MASH) arise, in part, from excessive free fatty acid flux from adipose tissue to the liver. Activin E (ActE, encoded by *INHBE*) suppresses adipocyte lipolysis through the type I receptor ALK7 (ACVR1C). Loss-of-function variants in *INHBE* and *ACVR1C* reduce waist-to-hip ratio in humans, yet genetic knockouts in mice produce insulin resistance and hepatic steatosis, suggesting discrepancies between human and mouse biology. We identify GDF3, a TGF-β superfamily ligand upregulated in obese adipose tissue, as the principal endogenous antagonist of ActE/ALK7 signaling. Human transcriptomic datasets reveal coordinated dysregulation: hepatic *INHBE* expression and circulating ActE protein are elevated in obesity, while adipose *ACVR1C* is downregulated and *GDF3* is reciprocally upregulated. Using ALK7-selective reporter assays, we show GDF3 inhibits ActE-driven SMAD2/3 signaling as a competitive antagonist rather than the weak agonist previously proposed. ActE suppressed beta-adrenergic-stimulated lipolysis in mouse and human adipocytes and primary human adipose tissue; GDF3 overexpression abolished this effect. In diet-induced obese mice, inducible *Gdf3* deletion reduced adipose lipolysis, resolved hepatic steatosis and fibrosis, and improved insulin sensitivity, benefits abolished by *Inhbe* knockdown, confirming dependence on ActE signaling. Predicted loss-of-function variants in *INHBE* show only nominal, WHR-dependent associations with type 2 diabetes risk, potentially confounded by hematological effects on HbA1c. *Gdf3* deficiency synergized with the clinical-stage anti-activin receptor antibody Bimagrumab to amplify fat-mass loss and improve glucose homeostasis in multiple MASH models. These findings establish GDF3 as an endogenous antagonist of ActE-ALK7 signaling and nominate GDF3 inhibition as a therapeutic strategy for MASLD/MASH.

## Introduction

Metabolic dysfunction-associated steatotic liver disease and steatohepatitis (MASLD, MASH) are rapidly growing global health concerns. The central driver of MASLD pathogenesis is dysregulated adipose tissue lipolysis, which typically accompanies obesity (*1*). Excessive free fatty acid flux from adipose tissue to the liver promotes hepatic steatosis, inflammation, and fibrosis, yet the mechanisms precipitating this dysregulated lipolysis remain incompletely understood.

While multiple signaling pathways may mediate the phenotypic effects of MASLD, activins and bone morphogenetic proteins (BMPs) have recently emerged as regulators of adipose tissue lipolysis (*2, 3*). The broader TGF-β superfamily comprises more than 30 secreted ligands, including TGF-βs, activins, BMPs, and growth differentiation factors (GDFs), that signal through heteromeric complexes of type I and type II serine/threonine kinase receptors to regulate diverse biological processes (*4*). Within this superfamily, Activin E (ActE), encoded by *INHBE*, is a liver-enriched hepatokine that signals through the type I receptor ALK7 (encoded by *ACVR1C*) in adipocytes, thereby suppressing lipolysis. Human genetic studies have implicated Activin E (*INHBE*) and ALK7 (*ACVR1C*) as mediators of waist-to-hip ratio (WHR) (*5–8*). Predicted loss-of-function variants in both genes reduce WHR, leading to the assumption that ActE deficiency protects against type 2 diabetes (T2D) (*6, 7*). Paradoxically, mouse knockouts of *Inhbe* or *Acvr1c* produce adverse metabolic phenotypes: unrestrained adipose lipolysis, insulin resistance, and hepatic steatosis, that directly contradict the protective effect inferred from human genetics (*9–14*). This discrepancy remains an important unresolved question as therapeutics targeting this pathway are developed.

We previously identified GDF3, a TGF-β superfamily ligand predominantly expressed in adipose tissue, as an activator of adipose lipolysis (*15*). GDF3 is a structural and functional outlier that has been shown to act as a BMP signaling antagonist and a weak activin-like signaling agonist through ALK7/ALK5, requiring either supraphysiological doses or the coreceptor CRIPTO, which is not expressed in adipocytes (*16–22*). Within adipose tissue, Gdf3 is controlled by PPARγ: thiazolidinediones suppress Gdf3 (*22*), partly by reducing PPARγ serine-273 phosphorylation, to link Gdf3 to insulin-sensitizing pharmacology (*22–25*). Overexpression of *Gdf3* renders even lean mice insulin resistant (*22*). Consistent with a clinical counterpart to this phenotype, *GDF3* expression showed a significant positive correlation with hemoglobin A1c (HbA1c), identifying glycemic dysregulation as its strongest clinical correlate (*26, 27*).

Here we show that GDF3 antagonizes ActE/ALK7 signaling, acting not as a weak agonist but as an endogenous inhibitor. In obese mice, inducible *Gdf3* deletion resolved hepatic steatosis. Gdf3 loss-of-function benefits were abolished by *Inhbe* knockdown, suggesting dependence on restored ActE signaling for lipolysis suppression. Additionally, we find that Gdf3 deficiency synergizes with the clinical-stage anti-activin receptor antibody Bimagrumab to amplify fat-mass loss and improve metabolic outcomes. These findings establish that GDF3 is the principal endogenous antagonist of ActE/ALK7 signaling and that its loss restores beneficial ActE activity to protect against MASH.

## Results

### Expression of ActE receptors in patients with obesity and liver disease

To identify the expression patterns of activin receptors in human physiology, we explored several published datasets and found that *ACVR1C* is the most highly expressed type I activin receptor in mature human adipocytes (**Fig. 1A**). All three type II receptors capable of binding activin proteins, *BMPR2*, *ACVR2A*, and *ACVR2B*, are expressed in mature human adipocytes (**Fig. 1B**). Within human omental adipose tissue, *ACVR1C* is selectively expressed in the adipocyte cell population and across all subtypes of mature adipocytes by single nucleus RNA-seq (*28*) (**Fig. 1C, D**). *ACVR1C* is more highly expressed in adipocytes from subcutaneous adipose tissue (SAT) of females versus males, and more highly expressed in visceral adipose tissue (VAT) compared to SAT (**Fig. 1E-1F**). Adipocyte expression decreases with obesity in SAT (**Fig. 1G**), consistent with reports across multiple studies demonstrating decreased expression of *ACVR1C* in the combined adipose tissue cell populations of SAT of obese females both with and without T2D (*28–30*) (**Fig. 1H)**. Expression is dynamic, as recovery of ACVR1C transcripts was observed through analysis of longitudinal weight loss intervention studies. Obese individuals who underwent successful weight reduction interventions showed restoration of *ACVR1C* expression to levels indistinguishable from those observed in lean controls (**Fig. 1I**). These data indicate that receptor downregulation is reversible and responsive to improvements in metabolic status. These publicly available data are in agreement with previously reported regulation of *ACVR1C* in mouse and human adipose tissue (*31–34*), consistently demonstrating an inverse correlation of *ACVR1C* expression with obesity and T2D. This regulation appears conserved between humans and mice as we observe a comparable expression pattern in adipose single-nucleus data from mice on a standard chow diet or high-fat diet (HFD) (*28*) (**Supplementary Fig. 1).**

**Figure 1.**
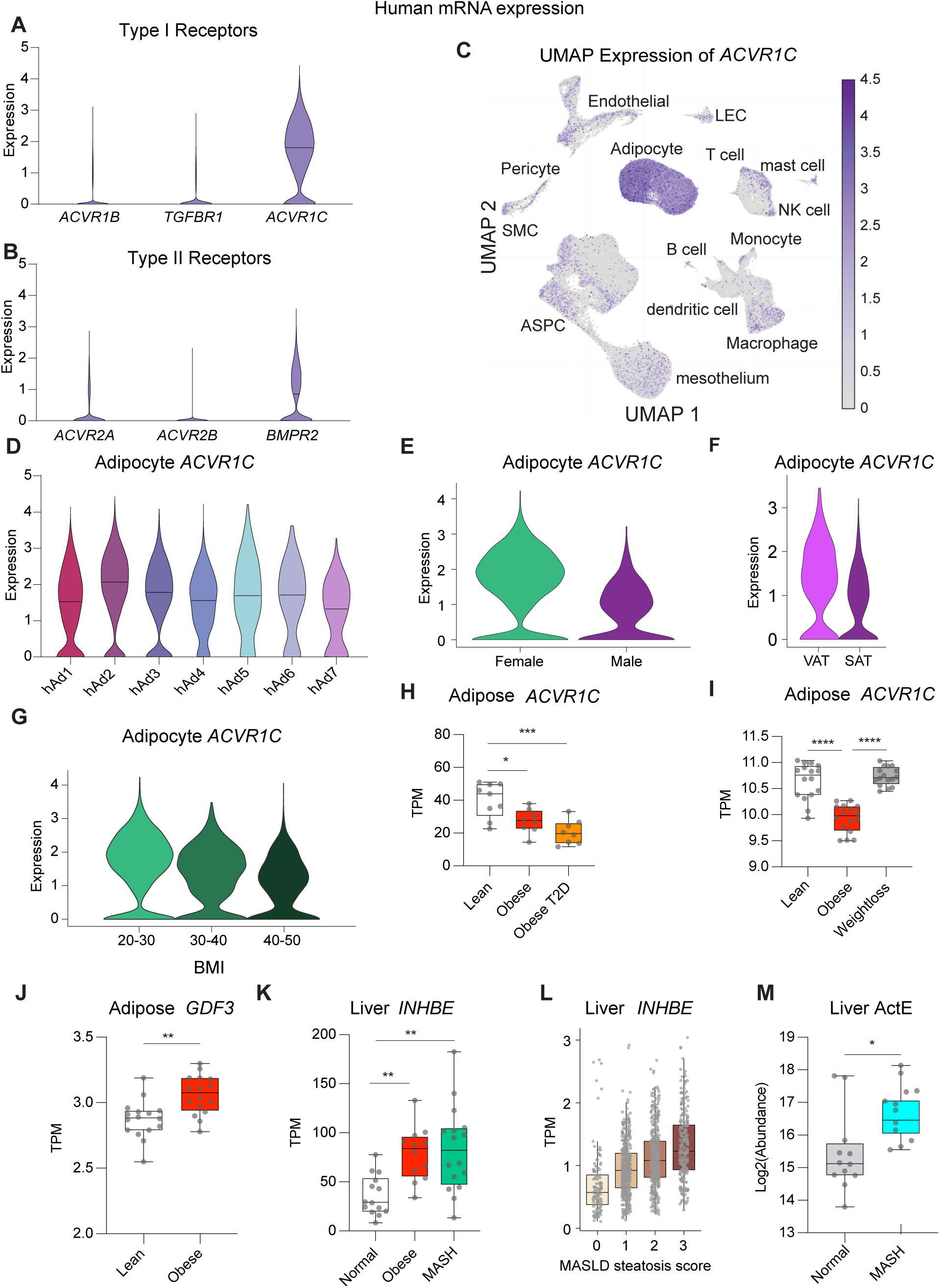
Activin receptor ALK7 and ligands are dysregulated in human obesity and MASH. **(A-F)** mRNA expression by single nucleus RNA-seq (*28*) collected from patients with body mass index (BMI) 20-30 kg/m^2^. **(A)** Expression of type I receptors (*ACVR1B*, *TGFBR1*, *ACVR1C*) and **(B)** type II receptors (*ACVR2A*, *ACVR2B*, *BMPR2*) in adipocytes. **(C)** UMAP representation of *ACVR1C* across adipose tissue (*28*). **(D)** *ACVR1C* expression distribution across previously identified adipocyte subpopulations (*28*). **(E)** Adipocyte *ACVR1C* expression in subcutaneous adipose tissue (SAT) from female and male patients. **(F)** Adipocyte *ACVR1C* expression SAT and visceral adipose tissue depots (VAT) from male patients (*28*). **(G)** *ACVR1C* expression with increasing BMI in SAT adipocytes from female patients (*28*). **(H)** Transcripts per million (TPM) of *ACVR1C* expression in adipose tissue across metabolic phenotypes: lean, obesity, and obesity with type 2 diabetes (T2D) (*83*). **(I)** TPM of *ACVR1C* expression dynamics in adipose tissue from lean, obese, and post-weight loss individuals (*84*). **(J)** TPM of *GDF3* in adipose tissue from lean versus obese individuals (*83*). **(K)** TPM of *INHBE* in liver tissue from healthy BMI, patients with obesity, and patients with MASH (*35*). **(L)** TPM of *INHBE* in liver tissue from healthy patients versus patients with increasing steatosis score with MASLD (n = 1,172) (*36, 37*). **(M)** Protein abundance of activin E in human livers from healthy and metabolic dysfunction-associated steatohepatitis (MASH) patients by proteomic determination (*85*). Data in (A–G) are shown as violin plots of expression distribution. Data in (H–M) are shown as box-and-whisker plots (median, interquartile range, min to max). P values determined by unpaired two-tailed t test for two-group comparisons (J, L, M) and one-way ANOVA with Tukey’s multiple comparisons test for comparisons of three or more groups (H, I, K).

### GDF3 and ActE upregulation accompanies ACVR1C downregulation in obesity

While adipose *ACVR1C* is downregulated with obesity, we identified reciprocal changes in expression of both ALK7 ligands ActE and GDF3. While *ACVR1C* is highly expressed in adipocytes, *GDF3* is expressed at levels too low for reliable detection by single nucleus RNA-seq analysis. However, the increased depth of bulk RNA-seq analysis revealed significant *GDF3* upregulation in adipose tissue samples from obese individuals compared to lean controls, consistent with previous findings with *Gdf3* elevation in mice on HFD (*19, 20, 22*) (**Fig. 1J, Supplemental Fig. 1E**). In previously reported findings, *INHBE* mRNA showed increased expression in patients with obesity and obesity-associated liver dysfunction (*5, 9*). Transcriptomic analysis of publicly available hepatic RNA sequencing datasets similarly demonstrated significantly increased *INHBE* mRNA expression in patients with obesity and MASH relative to lean controls (**Fig. 1K**) (*35*). This finding is consistent with the observed elevation of *INHBE* transcripts in MASLD liver relative to healthy controls (**Fig. 1L**) (*36, 37*) . In addition to increased expression, protein levels of ActE in liver tissue are elevated in patients with MASH (**Fig. 1M**). These data collectively suggest that obesity and liver disease are characterized by coordinated regulation of the INHBE-GDF3-ACVR1C signaling axis, with simultaneous receptor downregulation in adipose tissue and ligand upregulation.

### Weak association of loss-of-function variants in *ACVR1C* and *INHBE* with T2D in patients

In contrast to the highly significant (5 x 10^-8^) association between pLoF variants in *INHBE* (*5*) with decreased waist-hip ratio (WHR), the effects of these genes on the risk of developing T2D are less clear. To investigate whether there was a statistically significant decrease in the odds ratio of developing T2D, we examined associations with variants that are predicted to cause loss of protein function using data from UK Biobank, comprising whole-genome sequence data from ∼500,000 participants linked with deep phenotyping data (*39, 40*). In our dataset, there are 21 pLoF variants for *ACVR1C* in a total of 38 carriers. The most common variant, Chr2:157541153-G:A, is found in 12 patients and produces a premature stop codon at *ACVR1C* 388 and eliminates approximately one third of the intracellular kinase domain. Using methods analogous to studies of WHR, we modeled the association with T2D adjusted for age, sex and BMI—all of which affect *ACVR1C* mRNA expression levels. This base model (Model 1) did not reach nominal statistical significance for an association with reduced T2D risk (p=0.254). Model 2 includes an adjustment for WHR but also fails to reach statistical significance. We also considered the role of these genes in red blood cell formation as activin receptor modulation can alter erythropoiesis (*41*), and increased red blood cell counts can artificially lower values of HbA1c, a surrogate marker used in diagnosing T2D (*42*). In Model 3, we add an adjustment for reticulocyte percentage, a marker of immature red blood cells. None of these models using aggregated pLoF variants alone were sufficient to suggest a statistically significant protection from T2D. As these variants are rare, we examined the power of our model to detect a large effect size (OR=0.15) and observed an 80.5% power. That is, if losing an allele of *ACVR1C* does affect T2D, the effect size is likely to be less than a 6-fold reduced risk. By design, these analyses exclude the contribution of missense mutants in the receptor, as therapeutic strategies are not aiming to replicate specific receptor variants, but loss of receptor signaling.

Within the population studied, there are more pLoF variants in the *INHBE* gene with 44 variants in 1,017 carriers. The most common variant, Chr12:57456093-G:C is found in 655 patients and represents a splicing variant that impacts intron removal between the two exons of the *INHBE* gene. The base model produces a nominal, but not a genome-wide significance for protection from T2D (p=0.015) with an odds ratio of 0.714. When including WHR, the nominal significance is lost (p=0.060), suggesting that the protection from T2D is accounted for by the decreased WHR. As pLoF variants in *INHBE* are significantly associated with altered reticulocyte percentage (p=0.03), adjusting for red blood cell formation slightly attenuates the effect seen in the base model, suggesting that the effect on T2D may have a hematologic component (**Supplementary Table 1**).

### ActE inhibits adrenergic-stimulated lipolysis in human and mouse adipocytes

To understand whether there are differential effects of ActE between human and mouse cells on ALK7 signaling, we examined lipolytic regulation in immortalized adipocytes treated with ActE for 24 h and then stimulated with a beta-adrenergic agonist to measure the change in free fatty acid release. In mouse adipocytes, we observed that purified recombinant ActE suppresses beta adrenergic-stimulated lipolysis (**Fig. 2A**). In human adipocytes we observe ActE produces a ∼70% suppression of stimulated lipolysis at either 150 or 300 ng/ml ActE, but not at 75 ng/ml (**Fig. 2B**). Treatment with any of the doses of ActE did not affect mRNA expression of mature adipocyte markers *PPARG, ADIPOQ, FABP4, FASN,* or *LEP* indicating no changes in adipocyte differentiation after ActE treatment. In contrast, ActE at 150 or 300 ng/ml significantly lowered the expression of lipolytic genes *LIPE, PNPLA2*, and *ADRB3* when compared to untreated controls (**Supplemental Fig. 2A, 2B**). In surgically resected primary human adipose tissue slices from a patient with obesity, we find ActE can significantly suppress lipolysis at the highest dose tested (**Fig. 2C**).

**Figure 2.**
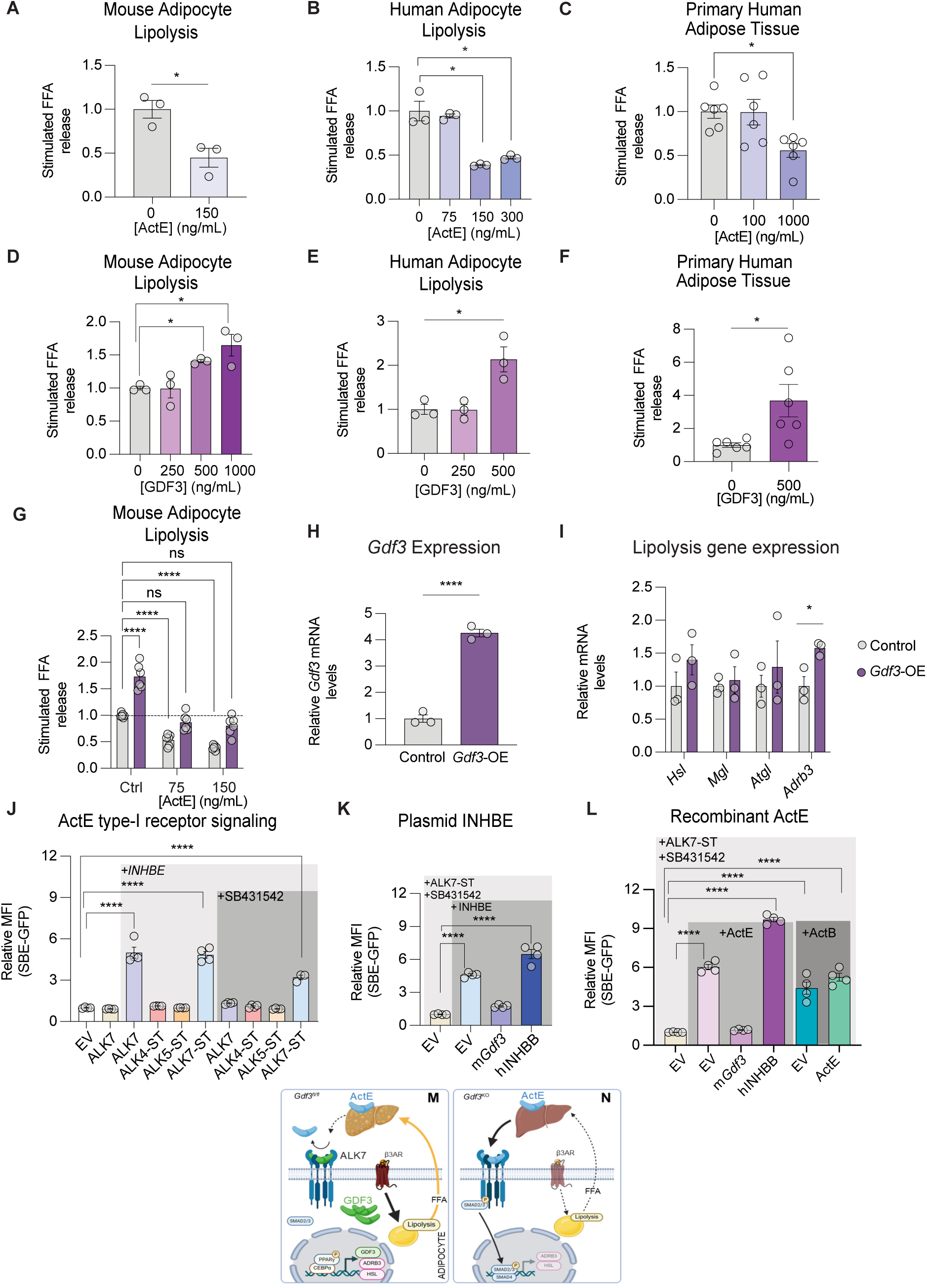
Activin E inhibits lipolysis via ALK7 signaling, antagonized by GDF3. **(A-G)** Fold induction of isoproterenol-stimulated free fatty acid release from cells treated for 24 h with ActE or GDF3. **(A)** Mouse immortalized adipocytes treated with recombinant ActE (150 ng/ml). **(B)** Immortalized human adipocytes treated with recombinant ActE (75, 150, 300 ng/ml). **(C)** Primary human adipose tissue from a patient with obesity treated with recombinant ActE (100, 1000 ng/ml). **(D)** Mouse immortalized adipocytes treated with recombinant GDF3 (250, 500, 1000 ng/ml) for 24 hours. **(E)** Immortalized human adipocytes treated with recombinant GDF3 (125, 250, 500 ng/ml). **(F**) Primary human adipose tissue from a patient with obesity treated with recombinant GDF3 (500 ng/ml). **(G)** Mouse immortalized adipocytes transduced with a control virus (Control) or virus overexpressing a constitutive *Gdf3* mRNA (*Gdf3*-OE). Lipolysis was measured either in the absence or presence of ActE (75 and 150 ng/ml). **(H, I**) Control and *Gdf3*-OE cells, mRNA of *Gdf3* or for lipolytic genes *Hsl*, *Mgl*, *Atgl*, and *Adrb3*. **(J - L)** Relative mean fluorescence intensity (MFI) of SMAD2/3 reporter binding element (SBE)-GFP in HEK293 cells transiently transfected with indicated plasmids (n = 4 biological replicates per condition; each n represents average relative MFI of approximately 2000 sorted live cells). **(J)** Reporter cells expressing plasmid-encoded *INHBE* and/or wild-type *ALK7* or SB431542-resistant *ALK4-ST*, *ALK5-ST*, or *ALK7-ST* were treated with or without SB431542, an inhibitor of wild type ALK4, ALK5, and ALK7. **(K)** Reporter cells expressing plasmid-encoded *ALK7-ST*, mouse *Gdf3*, or *INHBB* (encoding Activin B), treated with SB431542. **(L)** Reporter cells expressing plasmid-encoded *ALK7-ST*, *INHBE*, mouse *Gdf3*, or *INHBB* and treated with SB431542, recombinant activin E, or recombinant activin B as indicated. **(M)** Graphical representation of liver-adipocytes crosstalk in obesity. GDF3 expression is increase in obesity. Liver-derived ActE is antagonized by GDF3 bound to the ALK7 receptor complex. The absence of ActE/ALK7 signaling promotes transcription of ADRB3 enhancing catecholamine-driven lipolysis. Excess FFA in the serum are trafficked to the liver to promote steatohepatitis. **(N)** Gdf3 deficiency de-represses ActE/ALK7 signaling, leading to increased SMAD2/3 driven suppression of lipolysis. Reduced release of FFA prevents steatohepatitis. Data are mean ± SEM. P values determined by unpaired two-tailed t test for two-group comparisons (A, F, H), one-way ANOVA with Tukey’s multiple comparisons test for comparisons of three or more groups (B, C, D, E), and two-way ANOVA with Dunnett’s multiple comparisons test vs. the Ctrl: *Gdf3*-OE group for (G).

### GDF3 promotes adrenergic-stimulated lipolysis in human and mouse adipocytes

In contrast to the effect of ActE, GDF3 increases rates of adrenergic-stimulated lipolysis in both mouse and human and immortalized adipocytes (**Fig. 2D, 2E**). Similarly, in primary human adipose tissue slices, GDF3 can increase adipose tissue lipolysis (**Fig. 2F**). To determine whether this pro-lipolytic effect was mediated through ALK5 (*TGF*Β*R1*), an additional receptor for GDF3, we treated cells with the ALK5-selective inhibitors RepSox and galunisertib. The pro-lipolytic effect of GDF3 persisted in the presence of both inhibitors (**Supplementary Fig. 2C, 2D**), indicating that GDF3 promotes lipolysis through an ALK5-independent mechanism.

### GDF3 antagonizes ActE’s signaling and effects on lipolysis

While both GDF3 and ActE are annotated as ALK7 agonists (*15, 19, 43*), they produce opposite effects on adipose tissue lipolysis in vitro and in vivo (**Fig. 2A-2F)**. To understand the combined effects of GDF3 and ActE, we examined the effects of *Gdf3* overexpression in combination with recombinant ActE treatment. Increasing *Gdf3* expression increased stimulated lipolysis, similar to the effect of recombinant GDF3. This abolished the anti-lipolytic effect of exogenous ActE, demonstrating a dominant *Gdf3* effect (**Fig. 2G**); *Gdf3* overexpression was confirmed (**Fig. 2H**), and also increased *Adrb3* but did not affect *Hsl*, *Mgl*, or *Atgl* expression (**Fig. 2I**). We sought to better understand how ALK7 signaling is regulated by ActE and GDF3 ligands by creating an in vitro system that would signal exclusively through ALK7 without signaling contributions from either ALK4 or ALK5. We utilized HEK293 cells stably expressing a green fluorescent protein reporter downstream of a multimerized Smad-binding element that responds to SMAD2 and SMAD3 activation (SBE-GFP) (*44*). Plasmid-encoded *INHBE* can induce a 5-fold SMAD2/3 activation signal in cells co-transfected with wildtype *ALK7* (**Fig. 2J**) (*45*). However, while these cells have no endogenous ALK7 expression, they express endogenous ALK4/ACVR1B and ALK5/TGFBR1 (**Supplemental Fig. 2F**). To examine signaling exclusive to ALK7, we introduced the ST (serine-to-threonine) receptor system (*45, 46*). In the presence of the small molecule kinase inhibitor SB431542 (SB), all signaling through endogenous type I receptors (ALK4, ALK5, and ALK7) is abolished. Type I receptors with a serine to threonine mutation in the kinase domain’s ATP binding pocket are resistant to chemical inhibition by SB. In the absence of SB treatment, we detect no SMAD2/3 signal from ActE with ALK4-ST or ALK5-ST, but ActE strongly stimulates signaling through ALK7-ST. In the presence of the SB inhibitor, signaling through wildtype ALK7 is abolished and no signal through ALK4-ST or ALK5-ST is observed. ActE only activates SMAD2/3 signaling in the presence of ALK7-ST, confirming that ActE signals exclusively through ALK7 (*45, 47*) (**Fig. 2J**).

We next examined the effect of GDF3 on ActE signaling within the type I receptor-specific ALK7-ST/SB assay system. As before, ActE stimulates reporter activity at 5-fold greater levels than control cells. When *INHBE* and *Gdf3* are co-expressed, we observed that mouse GDF3 significantly decreases ActE-mediated ALK7-ST reporter activity (**Fig. 2K**). To ensure that expression of other TGF superfamily ligands did not universally impact ALK7 signaling, we examined the effect of an *INHBB* plasmid to express ActB. Together with ActE, ActB can increase reporter activity.

In our model, both GDF3 and ALK7 are co-expressed in adipocytes while liver-derived ActE arrives from the circulation. We tested this model for whether GDF3 could inhibit signaling from recombinant ActE as effectively as plasmid-encoded *INHBE*. We found recombinant ActE activated SMAD2/3 reporter activity to 6-fold above controls. ActE/ALK7-ST signaling was similarly inhibited by the transfection of plasmid-encoded *Gdf3*. However, co-expression of *INHBB* with *INHBE* further increased reporter activity (**Fig. 2L**). Alone, recombinant ActB activates ALK7 mediated reporter activity. In combination with ActE, recombinant ActB has similar reporter activity as ActE alone. These data suggest a model where liver-derived ActE signals through adipocyte ALK7/ACVR2A or ACVR2B to increase SMAD2/3 signaling to limit beta-adrenergic activation of lipolysis (**Fig. 2M**). GDF3 may function as an antagonist of ActE/ALK7 signaling when expressed in the same cell as the target receptors. Inhibition of ActE/ALK7 signaling leads to increased beta-adrenergic activated adipose tissue lipolysis, increased circulating FFA levels, and steatohepatitis (**Fig. 2N**).

### GDF3 inhibition of ActE promotes insulin resistance

To further investigate the opposing regulation of ALK7 by GDF3 and ActE, we measured rates of beta adrenergic-stimulated lipolysis in vivo. We examined mice with inducible whole-body *Gdf3* deletion after 8 weeks of HFD feeding (*Gdf3^KO^*) or littermate control mice (*Gdf3^fl/fl^*). We additionally treated *Gdf3^fl/fl^* or *Gdf3^KO^* mice with *Inhbe* siRNA to knock down ActE expression (*Inhbe^KD^ or Gdf3^KO^*::*Inhbe^KD^*) (**Fig. 3A**).

**Figure 3.**
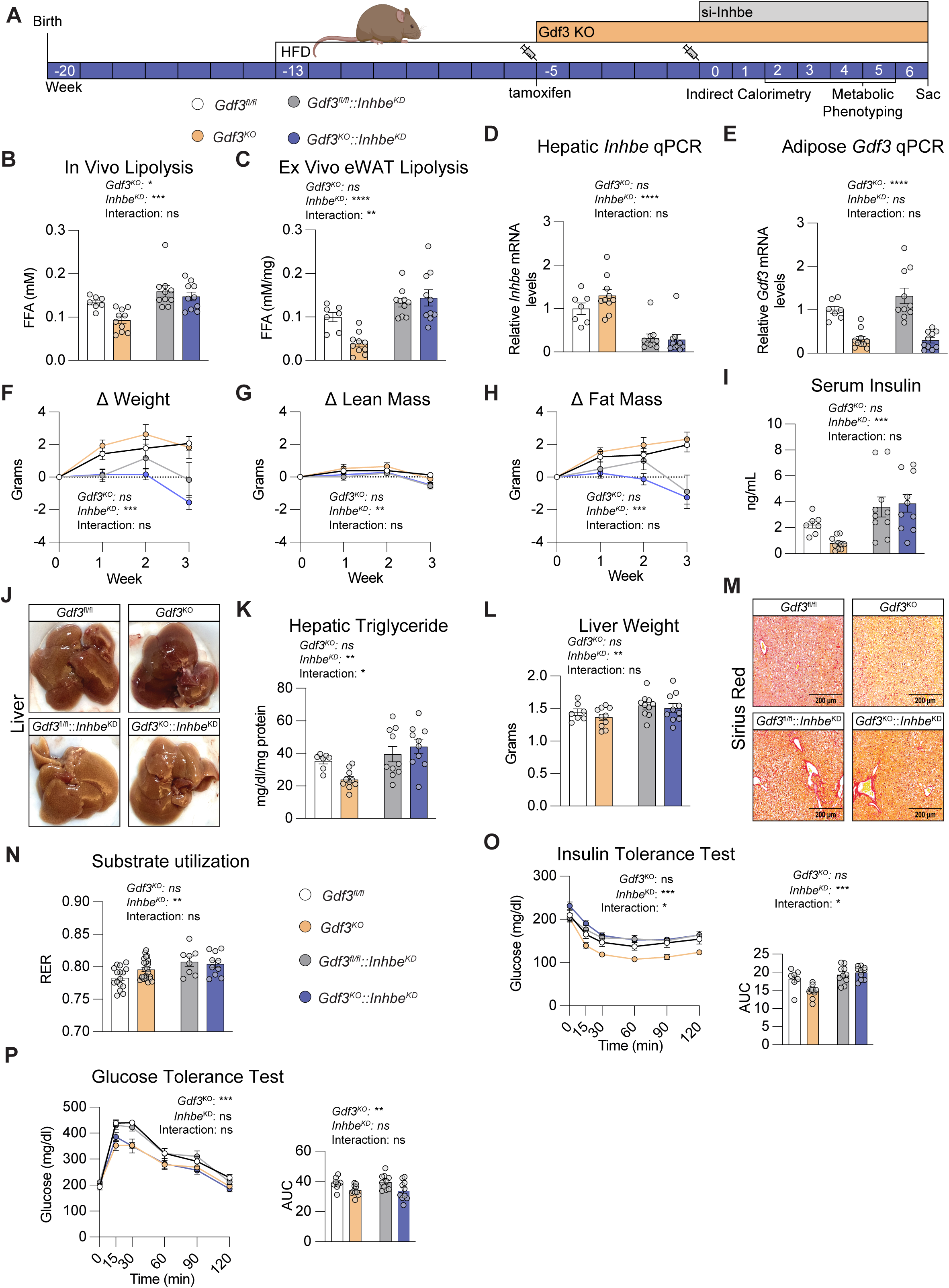
Activin E suppression drives excessive lipolysis and hepatic steatosis independent of GDF3 status. **(A)** Experimental design schematic. **(B)** In vivo lipolysis assessed by plasma non-esterified free fatty acid (FFA) concentrations 30 minutes following isoproterenol (ISO) administration. **(C)** Ex vivo lipolysis in epididymal white adipose tissue (eWAT) explants following ISO stimulation. **(D)** mRNA expression of *Inhbe* in mouse livers. **(E)** mRNA expression of *Gdf3* in mouse adipose tissue. **(F to H)** Percent change from baseline in total body weight **(F),** lean mass **(G),** and fat mass **(H)** over 3 weeks. **(I)** Fasting serum insulin concentrations. **(J)** Representative macroscopic liver images from *Gdf3^fl/fl^*, *Gdf3^KO^*, *Gdf3^fl/fl^*:*Inhbe*^KD^, and *Gdf3^KO^*::*Inhbe*^KD^ mice. **(K)** Total hepatic triglyceride content. **(L)** Liver weights. **(M)** Representative Sirius Red-stained liver sections. **(N)** Substrate utilization is represented as the respiratory exchange ratio (RER), measured by indirect calorimetry. **(O)** Insulin tolerance test (ITT) with area under the curve (AUC) quantification. **(P)** Glucose tolerance test (GTT) with AUC quantification. Indirect calorimetry measurements were performed 1 week following si*Inhbe* administration in ad libitum-fed mice. Data are mean ± SEM. Changes in body weight, lean mass, and fat mass over time (F to H) were analyzed by three-way repeated-measures ANOVA, with time as the within-subjects (matched) factor and Gdf3 genotype (*Gdf3^fl/fl^* versus *Gdf3^KO^*) and siInhbe (control versus *Inhbe*^KD^) as between-subjects factors. All other comparisons were analyzed by two-way ANOVA (Gdf3 genotype × siInhbe); the main-effect and interaction P values from this ANOVA are reported as text within each panel (Gdf3^KO^, Inhbe^KD^, interaction). For panels with a significant genotype × siInhbe interaction, Tukey’s multiple comparisons test was performed, and significant pairwise differences are indicated by brackets connecting the two groups compared. Asterisks therefore denote the pairwise comparison between the two groups that each bracket connects: *P < 0.05, **P < 0.01, ***P < 0.001, ****P < 0.0001; ns, not significant.

We measured FFA release both in vivo and in eWAT explants ex vivo. As previously described, *Gdf3^KO^*mice exhibit reduced rates of stimulated lipolysis compared to *Gdf3^fl/fl^* controls both in vivo and ex vivo (*15*). In the absence of ActE we observe a significant increase in beta-adrenergic receptor-stimulated lipolysis in vivo and ex vivo compared to *Gdf3^KO^* mice. However, reduced lipolysis seen with loss of *Gdf3* is abolished in the absence of ActE (**Fig. 3B, 3C)**. These findings are consistent with the biochemical observation that GDF3 blocks ActE/ALK7 signaling and the physiological effects of GDF3 require expression of ActE. Liver transcripts of *Inhbe* were lowered ∼70% by siRNA treatment irrespective of genotype (**Fig. 3D**). In the adipose tissue, *Gdf3* levels within both *Gdf3*^KO^ cohorts were lowered 70% (**Fig. 3E**).

### Decreased fat mass with increased serum insulin in *Inhbe*^KD^

Body weight and body fat increase over time for wildtype mice on the C57Bl/6 background on a 60% HFD (*48*). As in prior studies, inducible *Gdf3^KO^* did not alter body weight or body composition trajectories compared to *Gdf3^fl/fl^* controls. Conversely, *Inhbe^KD^* mice were resistant to weight gain on HFD and instead presented with significantly reduced fat mass, lean mass, and body weight relative to both *Gdf3^fl/fl^* and *Gdf3^KO^*controls (**Fig. 3F-3H)**. Measured levels of plasma insulin were non-significantly decreased with *Gdf3^KO^*but increased with *Inhbe^KD^* and *Gdf3^KO^*::*Inhbe^KD^* (**Fig. 3I**).

### GDF3 requires ActE to resolve hepatic steatosis

Given that dysregulated adipose lipolysis promotes fatty liver disease (*49*), we investigated whether GDF3 deficiency-driven reductions in lipolysis could attenuate hepatic steatosis in obese mice. When mice were maintained on a high-fat diet (HFD), control *Gdf3^fl/fl^*mice had livers with a pale, steatotic appearance (**Fig. 3J**). Livers from inducible whole-body *Gdf3^KO^* mice had a darker, healthier color associated with decreased liver fat. Conversely, loss of ActE signaling in both *Inhbe^KD^* and *Gdf3^KO^*::*Inhbe^KD^*mice was associated with markedly worsened hepatic steatosis, with increased triglycerides and liver weights evidenced by pale, lipid-laden livers that exceeded the severity observed in Gdf3^fl/fl^ controls (**Fig. 3J-3L**). While *Gdf3*^KO^ had a non-significant trend to lower hepatic triglycerides compared to control mice, this effect requires expression of ActE. *Gdf3*^KO^::*Inhbe*^KD^ mice had similar hepatic triglyceride levels as *Inhbe*^KD^ mice, with *Gdf3*^KO^::*Inhbe*^KD^ triglyceride rising significantly relative to *Gdf3*^KO^. Sirius red staining for liver fibrosis revealed a decrease in collagen deposition and fibrotic area for mice lacking *Gdf3* which was attenuated in the absence of ActE (**Fig. 3M**, **Supplementary Fig. 4A**). Only in *Gdf3^KO^* livers was the histological appearance consistent with decreased microsteatosis and fibrosis. To identify molecular drivers of these histological differences, expression profiling of adipose tissue revealed genes reciprocally regulated by *Inhbe^KD^* and *Gdf3^KO^*, our models of loss of ActE signaling and gain of ActE signaling. Transcripts showing altered regulation include *Pparg*, *Ppargc1a/Pgc-1a*, *Fasn*, *Acaca*/*Acc1*, and *Fto.* The regulated pathways in eWAT involved oxidative phosphorylation, lipolysis, fatty acid elongation, and branched chain amino acid degradation (**Supplemental Fig. 3A-3B**).

### GDF3 requires ActE for improvements in insulin tolerance

As we found that beneficial effects of *Gdf3^KO^* to decrease both liver steatosis and adipose lipolysis were lost in the absence of ActE signaling, we next investigated whether the systemic metabolic improvements seen with *Gdf3* deficiency were dependent on the expression of ActE. Indirect calorimetry revealed that *Inhbe^KD^* mice exhibited a significantly higher respiratory exchange ratio (RER) in the dark photoperiod, indicative of reduced fatty acid oxidation and increased rates of carbohydrate utilization (**Fig. 3N)**. In examining glucose metabolism, *Inhbe^KD^* impaired insulin tolerance, the decrease seen with *Gdf3^KO^ is* absent with *Inhbe^KD^*(**Fig. 3O**). Intriguingly, a distinct pattern of glucose tolerance was observed among the different treatment groups where *Gdf3* deficiency improved glucose tolerance independently of *Inhbe* expression (**Fig. 3P**). Together, these data demonstrate that the beneficial metabolic effects of *Gdf3* deficiency in obesity, reduced adipose tissue lipolysis, improved hepatic steatosis and fibrosis, and enhanced insulin sensitivity, are generally dependent on intact ActE signaling.

### GDF3 deficiency sustains metabolic protection during ACVR2A/2B blockade by Bimagrumab

Bimagrumab (BIMA) is a monoclonal antibody that neutralizes ACVR2A and ACVR2B, the type II receptors shared by myostatin, activins, and GDF11. In clinical trials, BIMA increases lean mass, reduces fat mass, and lowers HbA1c, with the greatest glycemic improvements observed when combined with the GLP-1 receptor agonist semaglutide (*50*). Because ACVR2A/2B are also required for ActE signaling, BIMA treatment would be expected to block ActE-mediated suppression of lipolysis and thereby promote hepatic steatosis, an effect that GDF3 deficiency might counteract by restoring ActE activity through an ACVR2-independent mechanism. We therefore asked whether combining GDF3 deficiency with BIMA would preserve metabolic benefits while still enabling the anabolic effects of activin blockade in skeletal muscle. To test this, we treated diet-induced obese *Gdf3^fl/fl^* and *Gdf3*^KO^ mice with BIMA for four weeks (**Fig. 4A**), confirming knockout of GDF3 in adipose tissue in *Gdf3*^KO^ mice (**Supplementary Fig. 4B**).

**Figure 4.**
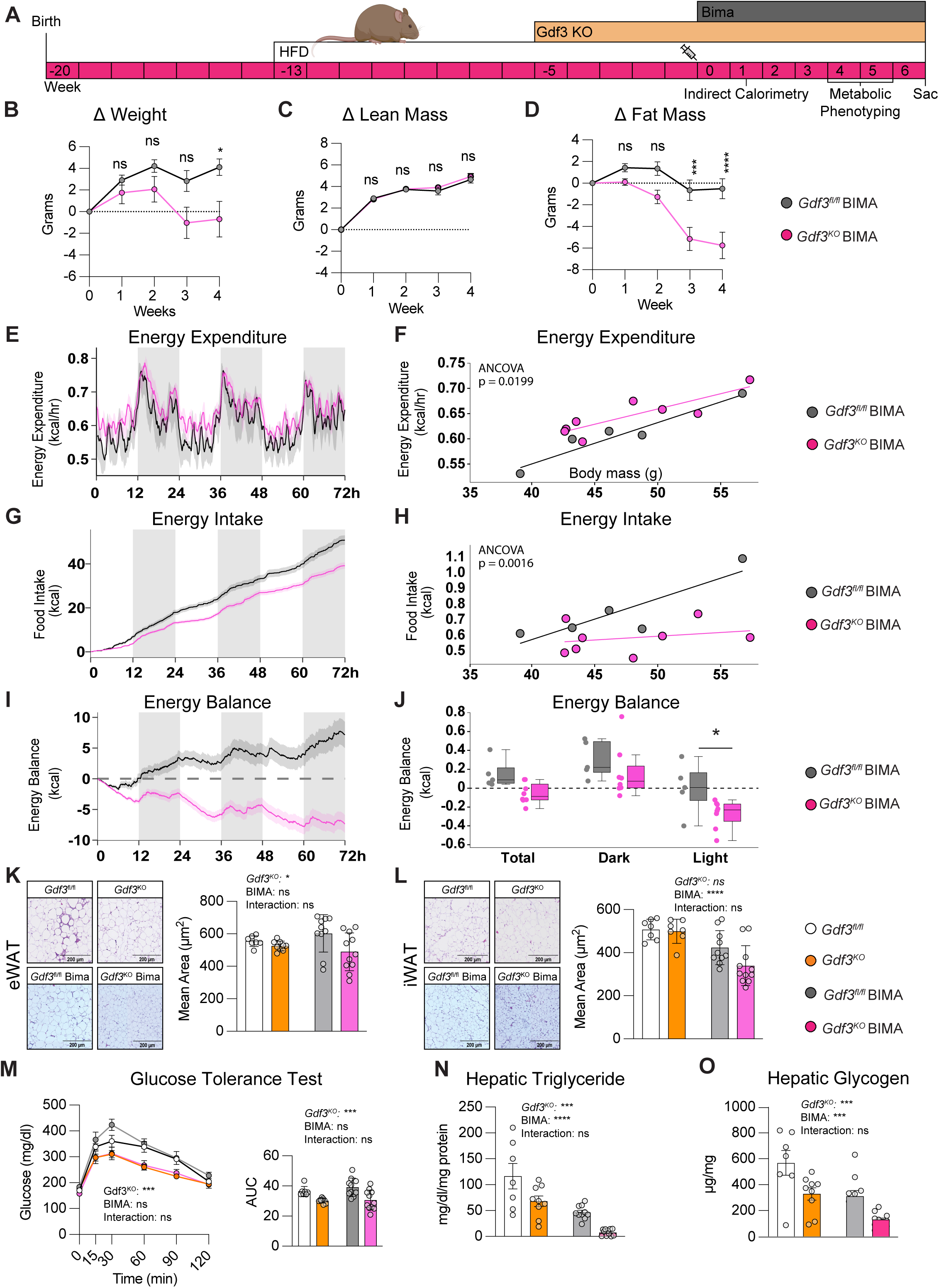
*Gdf3* deficiency with BIMA drives selective adipose loss and elevated energy expenditure. **(A)** Experimental design schematic. **(B to D)** Percent change from baseline in total body weight **(B),** lean mass **(C),** and fat mass **(D)** over 4 weeks of Bimagrumab (BIMA) treatment. **(E)** Energy expenditure (EE) measurements. **(F)** Linear regression analysis of EE versus body weight over 72 hours. **(G)** Cumulative food intake. **(H)** Linear regression of food intake versus body weight over 72 hours. **(I)** Total energy balance calculated from EE and energy intake over 72 hours. **(J)** Box plot representation of total energy balance (box, 25th–75th percentiles; whiskers, minimum–maximum; line, median). **(K)** Representative hematoxylin and eosin (H&E)-stained sections of epididymal white adipose tissue (eWAT) with quantification of mean adipocyte area. **(L)** Representative H&E-stained sections of inguinal white adipose tissue (iWAT) with quantification of mean adipocyte area. **(M)** Glucose tolerance test (GTT) with AUC quantification. **(N)** Total hepatic triglyceride content. **(O)** Hepatic glycogen content. Indirect calorimetry data were collected 2 weeks following the first BIMA administration in ad libitum-fed mice. Data are mean ± SEM. Changes in body weight, lean mass, and fat mass over time (B to D) were analyzed by two-way repeated-measures ANOVA, with time as the within-subjects (matched) factor and *Gdf3* genotype (*Gdf3*^fl/fl-BIMA^ versus *Gdf3*^KO-BIMA^) as the between-subjects factor, followed by Šídák’s multiple comparisons test; asterisks denote the *Gdf3*^fl/fl-BIMA^ versus *Gdf3*^KO-BIMA^ comparison at each time point. Energy expenditure (E, F) and food intake (G, H) were analyzed by analysis of covariance (ANCOVA) with body mass as the covariate. Energy balance (J) was compared between *Gdf3*^fl/fl-BIMA^ and *Gdf3*^KO-BIMA^. All remaining comparisons (K to O) were analyzed by two-way ANOVA (*Gdf3* genotype × Bimagrumab); main-effect and interaction *P* values are reported as text within each panel. For these panels, all pairwise comparisons among the four groups were made using Tukey’s multiple comparisons test, which corrects for multiple testing; significant differences are shown as brackets connecting the two groups compared, displayed only where a significant genotype × Bimagrumab interaction was present *P < 0.05, **P < 0.01, ***P < 0.001, ****P < 0.0001; ns, not significant.

Over the treatment period, BIMA-treated Gdf3^fl/fl^ mice gained approximately 4 g of body weight, whereas BIMA-treated Gdf3^KO^ mice lost approximately 1 g (**Fig. 4B**). Body composition analysis by quantitative NMR revealed that both genotypes gained a similar and substantial increase in lean mass (∼5 g), consistent with myostatin/activin blockade in skeletal muscle (**Fig. 4C**). However, fat mass responses diverged markedly with *Gdf3^fl/fl^* mice losing ∼1g and *Gdf3^KO^* mice losing ∼6 g (**Fig. 4D**). The pronounced fat loss in *Gdf3^KO^*mice was therefore sufficient to offset lean mass gains and produce net weight loss, in contrast to the net weight gain observed in *Gdf3^fl/fl^* controls.

### Enhanced fat loss in *Gdf3^KO^* is driven by increased energy expenditure and reduced food intake

To understand the mechanisms underlying this differential fat loss, we assessed energy balance by indirect calorimetry. Compared with BIMA-treated *Gdf3^fl/fl^* controls, BIMA-treated *Gdf3^KO^*mice displayed significantly higher energy expenditure during both light and dark photoperiods (**Fig. 4E, 4F**). Notably, *Gdf3^KO^* mice also consumed significantly less food during the light photoperiod (**Fig. 4G, 4H**); this reduction in food intake was not observed in *Gdf3^KO^* mice receiving vehicle alone, suggesting it is specific to the BIMA-treated context. When energy balance was calculated as the difference between energy intake and expenditure, BIMA-treated *Gdf3^fl/fl^* mice maintained a positive energy balance consistent with their net weight gain. In contrast, *Gdf3^KO^* were in a negative energy balance, explaining the observed weight loss (**Fig 4I, 4J**). Together, these data indicate that *Gdf3* deficiency amplifies fat loss during ACVR2 blockade by simultaneously increasing energy expenditure and reducing caloric intake.

### *Gdf3* knockout promotes selective fat loss during ACVR2 inhibition without redistribution of FFA to liver

The divergent fat mass responses between genotypes were confirmed histologically: BIMA-treated *Gdf3^KO^* mice showed significantly reduced mean adipocyte area in both epididymal and inguinal white adipose depots compared with BIMA-treated controls (**Fig. 4K, 4L**). A key concern with enhanced lipolysis is that mobilized fatty acids may be redirected to the liver rather than oxidized, worsening hepatic steatosis. To address this, we measured hepatic triglyceride and glycogen content. Both *Gdf3^KO^* and BIMA treatment individually reduced hepatic lipid accumulation; however, the combination produced the greatest reductions in hepatic triglycerides and glycogen, demonstrating that fat loss reflected true oxidative disposal rather than hepatic redistribution (**Fig. 4N, 4O**). These improvements were accompanied by reduced hepatic fibrosis markers and downregulated gluconeogenic gene expression (**Supplementary Fig. 4C, 4D**), consistent with improved hepatic metabolic function in the absence of GDF3.

Previous reports have noted that BIMA impairs glucose metabolism in mice (*51, 52*). In this model, BIMA did not significantly worsen glucose tolerance in *Gdf3^fl/fl^* mice. Strikingly, *Gdf3* deficiency improved glucose tolerance beyond even in the presence of BIMA, suggesting that the glucoregulatory benefit of GDF3 loss is independent of ACVR2A/2B signaling (**Fig. 4M**).

### *Gdf3* deficiency prevents Western diet-induced weight gain and fat accumulation

Preclinical studies of metabolic dysfunction-associated steatotic liver disease (MASLD) in rodents indicate that high-fat diet (HFD) models inadequately recapitulate the progression from fatty liver to cirrhosis observed in humans. Feeding a Western diet at thermoneutrality provides a more physiologically relevant model that better reflects human disease pathology (*53*). To further investigate the metabolic effects of Bimagrumab (BIMA) in the context of *Gdf3* deficiency in conditions that drive hepatic metabolic dysfunction, we employed a thermoneutral Western diet (**Fig. 5A**). Four experimental groups were established: Control (*Gdf3^fl/fl^*), knockout (*Gdf3^KO^*), BIMA-treated controls (*Gdf3^fl/fl^*^-BIMA^) and BIMA-treated *Gdf3* knockouts (*Gdf3^KO^*^-BIMA^). We confirmed knockout of *Gdf3* in adipose tissue in both vehicle and BIMA-treated groups *Gdf3^KO^* mice (**Supplementary Fig. 5A**). Similar to our previous observations with HFD, on Western diet, only mice receiving BIMA treatment in the absence of GDF3 displayed a significant weight loss effect, losing 12% of total body weight (∼6g) (**Fig. 5B**). While BIMA increased lean mass in *Gdf3^fl/fl^* and *Gdf3^KO^* groups without genotype-dependent differences (**Fig. 5C**), *Gdf3^KO^*^-BIMA^ mice exhibited significantly greater fat mass reduction than all other groups (**Fig. 5D**).

**Figure 5.**
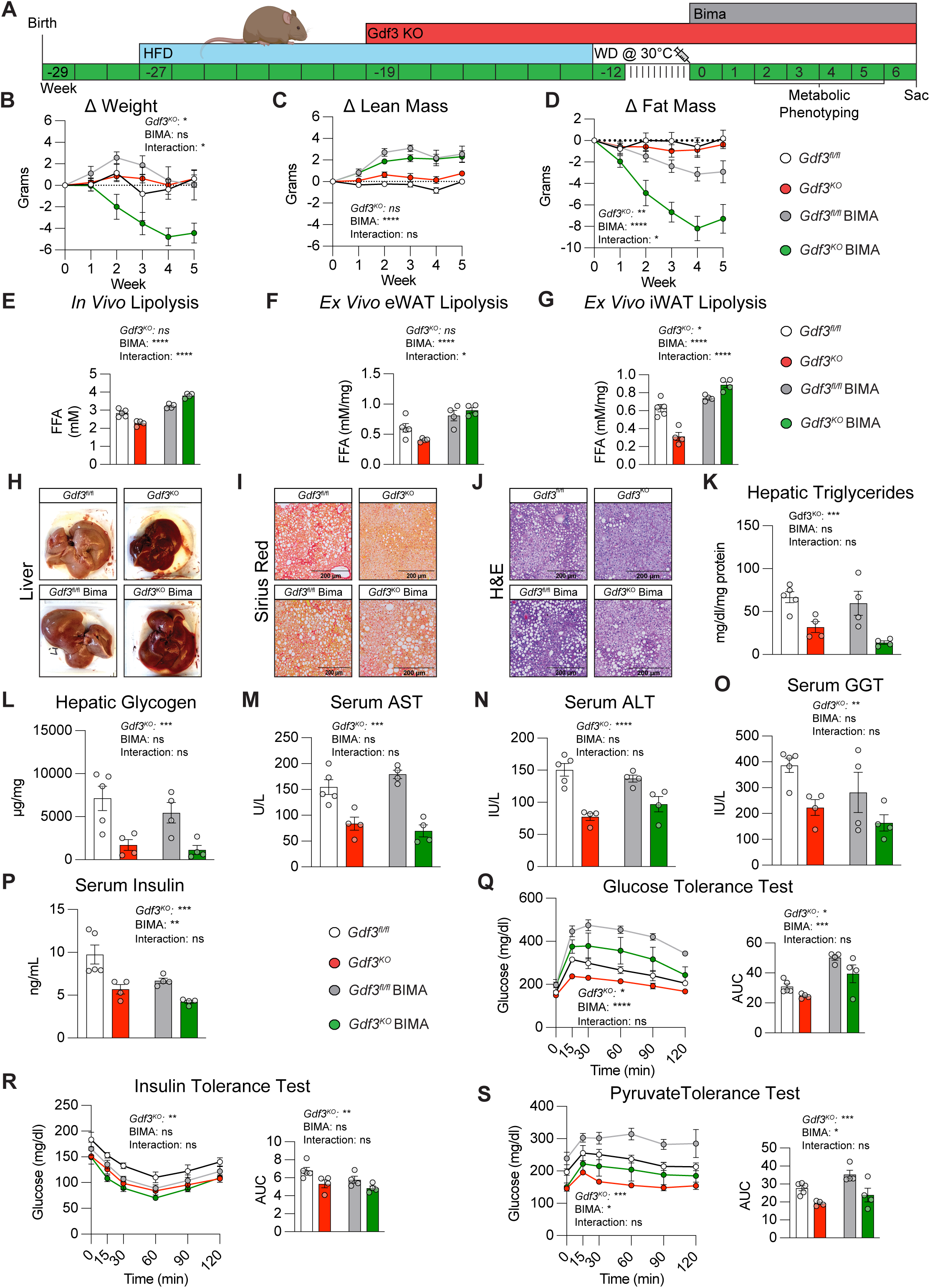
Combined *Gdf3* deletion and BIMA prevent diet-induced MASLD under thermoneutral conditions. **(A)** Experimental design schematic. **(B to D)** Percent change from baseline in total body weight **(B),** lean mass **(C),** and fat mass **(D)** over 5 weeks (2 weeks during BIMA treatment followed by 3 weeks post-treatment monitoring). **(E)** In vivo lipolysis assessed by plasma non-esterified fatty acid (NEFA) concentrations 30 minutes following isoproterenol (ISO) administration. **(F and G)** Ex vivo free fatty acid (FFA) release from eWAT **(F)** and iWAT **(G)** explants following isoproterenol stimulation. **(H)** Representative macroscopic liver images. **(I)** Representative Sirius Red-stained liver sections. **(J)** Representative H&E-stained liver sections. **(K)** Hepatic triglyceride content. **(L)** Hepatic glycogen content. **(M to O)** Serum aspartate aminotransferase (AST) **(M),** alanine aminotransferase (ALT) **(N),** and gamma-glutamyl transferase (GGT) **(O)** concentrations. **(P)** Fasting serum insulin concentrations. **(Q)** Glucose tolerance test (GTT) with AUC quantification. **(R)** Insulin tolerance test (ITT) with AUC quantification. **(S)** Pyruvate tolerance test (PTT) with AUC quantification. Data are mean ± SEM. Changes in body weight, lean mass, and fat mass over time (B to D) were analyzed by three-way repeated-measures ANOVA, with time as the within-subjects (matched) factor and *Gdf3* genotype (*Gdf3*^fl/fl^ versus *Gdf3*^KO^) and Bimagrumab (vehicle versus BIMA) as between-subjects factors; main-effect and interaction *P* values are reported as text within each panel. All other comparisons (E to S) were analyzed by two-way ANOVA (*Gdf3* genotype × Bimagrumab); main-effect and interaction *P* values are likewise reported as text within each panel. For these panels, all pairwise comparisons among the four groups (*Gdf3*^fl/fl^, *Gdf3*^KO^, *Gdf3^fl/fl^*^-BIMA^, *Gdf3*^KO-BIMA^) were made using Tukey’s multiple comparisons test, which corrects for multiple testing; significant differences are shown as brackets connecting the two groups compared, displayed only where a significant genotype × Bimagrumab interaction was present. *P < 0.05, **P < 0.01, ***P < 0.001, ****P < 0.0001; ns, not significant.

### Enhanced lipolytic activity underlies adipose tissue remodeling in *Gdf3*-deficient mice

To confirm the adipose mechanisms underlying the pronounced fat loss observed in BIMA-treated *Gdf3^KO^* mice, we assessed both in vivo and ex vivo lipolytic activity. BIMA treatment significantly increased circulating free fatty acid levels in vivo compared to control groups. These data are consistent with enhanced lipolytic activity due to inhibition of ActE signaling. Notably, *Gdf3^KO^* mice treated with BIMA exhibited more pronounced free fatty acid release relative to BIMA-treated controls, while *Gdf3^KO^* mice receiving vehicle showed significantly reduced free fatty acid levels, consistent with our previous HFD findings (**Fig. 5E**).

Ex vivo lipolysis assays using isolated adipose tissue provided mechanistic insights into BIMA’s effects in a more controlled environment. Both inguinal (iWAT) and epididymal (eWAT) white adipose tissues from BIMA-treated mice demonstrated significantly greater free fatty acid release compared to vehicle-treated controls (**Fig. 5F, 5G**). In iWAT from *Gdf3^KO^* mice, BIMA treatment produced a considerable enhancement in free fatty acid release compared to BIMA-treated controls, while *Gdf3^KO^* mice receiving vehicle showed markedly reduced lipolytic activity. Basal free fatty acid levels remained unchanged across all experimental conditions, indicating that the observed differences reflect treatment-specific alterations in lipolytic capacity rather than baseline metabolic variations. In cultured mouse adipocytes, BIMA treatment increased rates of lipolysis at all doses tested (**Supplemental Fig. 5B**). In primary human adipose tissue slices, BIMA also produced increased stimulated adipose tissue lipolysis (**Supplemental Fig. 5C)**. These effects are consistent with inhibition of ALK7 signaling by BIMA to increase adipose tissue lipolysis.

### *Gdf3* deficiency ameliorates Western diet-induced hepatic steatosis and fibrosis

Gross morphological examination revealed that livers from *Gdf3^KO^*mice exhibited reduced hepatic fat accumulation regardless of BIMA treatment status (**Fig. 5H**). Histological analysis using Sirius red staining demonstrated significantly lower collagen deposition and reduced hepatic fibrotic area in *Gdf3^KO^* mice with or without BIMA treatment (**Fig. 5I, Supplementary Fig. 5D**), accompanied by marked decreases in fibrosis and extracellular matrix markers (**Supplementary Fig. 5E, 5F**) .Further visualization by hematoxylin and eosin staining confirmed reduced hepatic lipid accumulation in Gdf3^KO^ mice regardless of BIMA treatment (**Fig. 5J**).

Biochemical analysis of liver tissue revealed that *Gdf3^KO^*mice showed significantly lower hepatic triglyceride and glycogen levels compared to controls, irrespective of BIMA treatment (**Fig. 5K, 5L**). Given that Western diet feeding at thermoneutrality serves as a model of hepatotoxicity, we assessed liver function markers and found a consistent pattern where *Gdf3^KO^* mice exhibited significantly reduced serum levels of alanine aminotransferase (ALT), aspartate aminotransferase (AST), and gamma-glutamyl transferase (GGT), independent of treatment status (**Fig. 5M-5O**).

### *Gdf3* deficiency preserves glucose homeostasis despite Bimagrumab-induced metabolic stress

After our previous observations that *Gdf3^KO^* improved systemic metabolism, we again assayed serum levels and markers which are dysregulated in obesity. Serum insulin measurements revealed that BIMA treatment alone, as well as *Gdf3^KO^* mice receiving PBS or BIMA, had significantly lower insulin levels than vehicle-treated *Gdf3^fl/fl^*controls (**Fig. 5P**).

We next evaluated glucose metabolism in all groups, which revealed distinct patterns of improvement divergent from those seen in HFD alone. As expected, glucose tolerance testing demonstrated that *Gdf3^KO^*mice exhibited improved glucose clearance compared to genotype-matched controls across both treatment conditions (**Fig. 5Q**). In PBS-treated groups, these findings parallel our previous HFD observations where *Gdf3* deficiency improved glucose tolerance despite comparable body composition due to reduced adipose tissue lipolysis. Notably, while both groups of BIMA-treated mice showed glucose intolerance relative to vehicle controls, Gdf3^KO^ mice maintained superior glucose tolerance compared to PBS or BIMA controls regardless of treatment status. Insulin sensitivity assessments confirmed that *Gdf3^KO^*mice demonstrated significant improvements in insulin tolerance with or without BIMA treatment (**Fig. 5R**). The positive effect of *Gdf3^KO^* was indistinguishable from BIMA treated *Gdf3^fl/fl^* controls which also displayed an equally improved insulin tolerance. Pyruvate tolerance testing, used to assess hepatic glucose production, revealed elevated glucose responses in BIMA-treated mice compared to vehicle controls (**Fig. 5S**). Notably, *Gdf3^KO^* mice exhibited significantly lower fasting glucose levels and attenuated pyruvate-induced glucose production compared to their respective controls, consistent with downregulation of hepatic gluconeogenesis markers (**Supplementary Fig. 5G**).

Our results align with recent studies demonstrating that BIMA or ACVR2B-Fc administration increases blood glucose levels and affects glucose regulation in mice (*51, 54–57*), highlighting the complex interplay between activin signaling and metabolic homeostasis. These findings underscore the differential roles of GDF3 in the absence of ActE signaling, and with ACVR2A/2B receptor blockade with BIMA. Loss of GDF3 has minimal impact without ActE expression, but *Gdf3^KO^* can continue to mediate a metabolic improvement when combined with BIMA.

## Discussion

Our understanding of activin biology and metabolism is at a crossroads. Human genetic association studies suggest that loss of ActE/ALK7 is protective in reducing diabetes risk. Clinical studies reveal elevated *INHBE* expression and ActE protein in the liver of individuals with obesity. This pattern and the association of *INHBE* loss-of-function mutants with lowered WHR have led to the interpretation that ActE is metabolically harmful, implying that reducing ActE signaling would be beneficial. However in vivo mechanistic studies from rodents and in vitro studies with mouse and human cells, including this study, demonstrate the opposite conclusion. We find that ActE signaling is protective in preserving metabolic health. In vivo, we find that disruption of ActE, the type I receptor ALK7, or the type II receptors ACVR2A/2B all increase adipose tissue lipolysis, promote metabolic dysfunction and accelerate the development of liver steatosis. How do we explain the apparent discrepancy between the suggestion of protective effects on T2D from human pLoF genetics and a phenotype of metabolic impairment in mouse KO studies?

In this study, we find that ActE functions identically in cultured adipocytes from mouse and human and in primary human adipose tissue; ActE promotes a beneficial decrease in lipolysis by downregulation of catecholamine receptor expression. As there is no obvious phenotypic difference between signaling in human and mouse cells, we may wish to carefully examine human interventions that propose to impede ALK7 or ActE signaling in vivo for adverse effects on liver health.

In addition, we propose a re-evaluation of the human genetic association studies with regard to protection from T2D. Variants in both *ACVR1C* and *INHBE* are significantly associated with decreased WHR, a phenomenon that *could* translate into protection from T2D. Yet the significance of pLoF variants in these genes with protection from T2D is unpersuasive. As a parallel example, pLoF variants in the insulin receptor gene (*INSR*) are also strongly associated with decreased WHR but increase the risk for T2D (*5, 8*). The explanation for this result is that insulin signaling generates a powerful anti-lipolytic signal. The effect seen with loss of *INSR*, *ACVR1C*, or *INHBE* is increased adipose tissue lipolysis preferentially from visceral depots and serves to decrease WHR and increase circulating FFA. Here loss of *INSR* is not protective from T2D despite the decreased WHR. That is, pathological genetic variants that reduce WHR may not produce the same benefit as lower WHR seen in epidemiological studies. We posit that unrestrained adipose tissue lipolysis is adverse for metabolic health without changes in body weight. However, with the anabolic effects of myostatin signaling inhibition, the increased FFA released with lipolysis can safely fuel muscle growth.

Activin proteins and growth differentiation factors have multiple physiological roles including to inhibiting differentiation. This is seen for myostatin (GDF8) in muscle, GDF3 in embryonic stem cells, and activins in erythropoiesis (*58–60*). Patients with anemia benefit from therapeutic inhibitors of activin signaling. Luspatercept and Sotatercept are FDA approved ligand traps, Fc-fusions of type II activin receptors which increase red blood cell formation (*41*). Consistent with this, *INHBE* pLoF variants are associated with significantly altered reticulocyte percentage (*5*). In addition, genetic association studies find markers for increased red blood cell formation in patients with predicted loss-of-function variation in *INHBE* and *ACVR1C* genes. The interpretation of HbA1c as the relative amount of glycated hemoglobin assumes a normal steady-state production and clearance rate of erythrocytes. For example, patients with both T2D diabetes and hemolytic anemia in which breakdown and clearance of erythrocytes is accelerated, will present with artificially reduced HbA1c levels, inconsistent with glycemic traits (*42*). Given the altered hematological parameters observed in patients with *ACVR1C* and *INHBE* variants, further research is required to understand whether loss of *ACVR1C* or *INHBE* confers *bona fide* protection from T2D. The lack of a strong human genetic signature for GDF3 loss-of-function may be attributable to the requirement of GDF3 in development. Studies have noted that embryonic loss of *Gdf3* results in lethality for ∼1 in 3 mice (*19, 21, 61*). In contrast, GDF3 appears to be dispensable and likely maladaptive in mature animals, and its inhibition enables the beneficial effects of preserved ActE activity (*62*). These results suggest that modulating GDF3 rather than direct ALK7/ActE blockade may avoid these adverse metabolic trade-offs.

This study clarifies why elevated ActE levels seen in obesity are not associated with a beneficial metabolic profile. We establish GDF3 as an endogenous obesity-induced antagonist of the liver-adipose ActE-ALK7 signaling axis. Our model suggests that elevated GDF3 levels in obesity contribute to metabolic dysfunction by disrupting ActE’s suppression of adipose tissue lipolysis (*63*). We show that *Gdf3* deficiency in obese adult mice provides comprehensive metabolic protection by reducing hepatic steatosis and fibrosis, improving glucose homeostasis, and enhancing fat mobilization in a manner dependent on ActE expression. These findings resolve the paradox of why ActE fails to suppress lipolysis effectively in obesity despite being markedly upregulated (*9*). Our studies identify GDF3 as a potential therapeutic target for MASLD and metabolic disease.

ALK7 regulation of lipolysis is a feature shared by all adipocytes in both mice and humans (*64*). ALK7 is more highly expressed in females compared to males. This finding may provide insight into the greater magnitude effect and statistical association of *ACVR1C* pLoF variants on WHR in females (*8*). The greater expression in VAT compared to SAT may also contribute to the greater rates of lipolysis in VAT vs SAT in obesity (*65*). Analysis of human transcriptomic datasets reveal the coordinated dysregulation of the ActE-ALK7 axis in obesity: hepatic *INHBE* and circulating ActE protein are elevated, adipose *ACVR1C* (ALK7) is downregulated, and adipose *GDF3* is upregulated (*9*). This pattern suggests a maladaptive feedforward loop. As obesity progresses and hepatic lipids accumulate, the liver increases ActE production as a compensatory attempt to suppress adipose lipolysis and to limit further FFA influx. However, simultaneous upregulation of GDF3 and downregulation of ALK7 in adipose tissue prevent ActE from exerting its anti-lipolytic effects. The resulting sustained lipolysis delivers excess FFAs to the liver, perpetuating steatosis, inflammation, and fibrosis.

These data suggest an updated model where adipose GDF3 functions as a competitive antagonist of ActE for shared receptors, rather than as a weak ALK7 agonist (*15, 16, 19, 66, 67*). In the adipocyte context, GDF3 is an antagonist of an inhibitory signal for lipolysis, which explains the previous phenotypic attribution as a pro-lipolytic agonist. Although recombinant GDF3 can activate a phospho-SMAD2/3 signal in ALK7-dependent reporter assays, its potency as an agonist is 100-fold weaker than that of canonical TGF-β ligands (*18–20*). More importantly, high concentrations of recombinant GDF3 are capable of stimulating adipose tissue lipolysis in murine cellular and explant models (*15*), the opposite effect of ActE-ALK7 activation to suppress lipolysis (*10, 12*). From our in vivo and in vitro data, we propose that the principal effect of GDF3 in obesity and metabolic disease is its antagonism of ActE. By competing for shared mature adipocyte receptors with insufficient intrinsic agonist activity to activate downstream SMAD2/3 signaling, GDF3 may prevent ActE from signaling without itself inducing a signal. This mechanism effectively disrupts the liver-adipose feedback loop that normally restrains FFA mobilization by desensitizing the adipose to ActE signaling.

While we had initially hypothesized that metabolic improvements seen with *Gdf3^KO^* would be lost if ActE could not signal through ACVR2A/2B due to neutralization with BIMA, instead, we found that despite increased rates of stimulated lipolysis, glucose regulatory and hepatoprotective effects of *Gdf3^KO^*were maintained. Unexpectedly, we also observed a synergistic fat loss effect of BIMA-induced activin signaling blockade in the absence of GDF3. Although both GDF3 and myostatin can signal through ALK5, *Gdf3^KO^* mice show no alterations in muscle mass under basal conditions or during neutralization with BIMA, demonstrating that GDF3 does not compensate for myostatin loss in muscle growth regulation. Instead, GDF3 deficiency potentiates fat loss during ACVR2 inhibition, resulting in selective reduction of fat mass without affecting lean mass gains. This suggests a unique metabolic role for GDF3 in controlling adipose energy release, distinct from myostatin’s muscle-centric actions, and in improving liver and whole-body metabolism during anabolic states. These findings suggest that combining GDF3 inhibition with anabolic agents (the myostatin signaling inhibitor BIMA) could provide synergistic benefits, including muscle hypertrophy, enhanced fat mobilization, and metabolic improvements.

When *Gdf3^KO^* or *GDF3^fl/fl^* mice were challenged with western diet and housed at thermoneutrality, the fat loss effect of BIMA treatment in *Gdf3^KO^*was maintained. Consistent with previous observations, BIMA treatment alone was sufficient to produce a slight decrease in fat mass but was significantly lower than the combination of BIMA and *Gdf3^KO^*. Despite elevated in vivo and ex vivo stimulated lipolysis rates with BIMA treatment, *Gdf3^KO^* prevented hepatic lipid accumulation and the resulting fibrosis and inflammation indicative of a MASH-like phenotype. *Gdf3^KO^* improved the impaired glucose tolerance and attenuated the exaggerated pyruvate response observed in BIMA-treated mice. Together, these data suggest that even in a model of MASH-associated liver toxicity, *Gdf3^KO^* can improve metabolic health. Although BIMA likely inhibits ActE/ALK7 signaling in adipose tissue, the sustained benefit of *Gdf3^KO^* indicates additional mechanisms. It is possible that the increased energetic demand of muscle growth may compensate for elevated lipolysis in the adipose tissue and prevent FFA accumulation in the liver (*68*). Future studies will be needed to determine whether increased adipose tissue lipolysis combined with a weight-loss regimen could promote selective loss of fat mass and preservation of muscle mass. GDF3 inhibition is predicted to be beneficial for whole-body glucose metabolism and resolution of steatohepatitis in weight-neutral or weight-gain conditions. GDF3 inhibition in relative negative energy balance is also likely to promote selective loss of fat mass as seen with the *Gdf3^KO^* mice treated with BIMA.

## Limitations

This model allows us to integrate and re-evaluate previous reports of GDF3 and ActE biology. GDF3 requires ALK7 for SMAD2/3 regulation, as shown in ALK7 KO cells (*12*). Beneficial effects of GDF3 require intact ALK7 signaling, as shown with transplantation of *Gdf3^KO^* bone marrow into ALK7-deficient mice (*20*). The only finding our model does not account for is the observation by Bu et al. that GDF3 decreases lipolysis in vitro (*20*). The observed differences may be attributable to differences in the adipocytes derived from the TSOD mouse strain in their studies that spontaneously develop diabetes (*69–71*). Within the context of broader activin and SMAD2/3 signaling, the function of GDF3 remains unclear. Many studies have indicated that myostatin, Activin B, or the secreted antagonist FSTL3 have strong effects on adipocyte development and systemic metabolism within the context of obesity (*72–79*). As inhibition of activin signaling has shown clinical relevance for increasing lean mass, decreasing fat mass, and improving systemic insulin tolerance in HFD, we are interested in the capacity of GDF3 to inhibit additional ligands within these contexts (*51, 80–82*). While our current model of GDF3 antagonism within the adipocyte supports direct competition for receptors between GDF3 and ActE, it does not exclude other models of noncompetitive antagonism. Additionally, one possible explanation for the differences between human and mouse biology may relate to the subtle differences in protein structures between species, with ACVR2A/B having >99% protein identity, ALK7 with 93% identity, but GDF3 is only 71% identical.

We propose an updated model of adipose-liver metabolic regulation in which GDF3 acts as the key endogenous antagonist of the beneficial actions of ActE. In lean, metabolically healthy states, very low GDF3 levels do not block ActE, allowing effective regulation of adipose lipolysis by catecholamines. Furthermore, GDF3 deficiency in lean mice has no effect on glucose homeostasis (*22*). In obesity, increased GDF3 antagonizes ActE, disrupting this protective feedback loop and promoting sustained lipolysis, hepatic lipid accumulation, and metabolic dysfunction. Genetic deletion of GDF3 restores ActE efficacy, reduces lipolysis, resolves hepatic steatosis and fibrosis, and improves glucose homeostasis across multiple models. These findings nominate GDF3 inhibition as a novel therapeutic strategy for obesity-induced MASLD/MASH and provide a mechanistic framework for optimizing combination therapies targeting the TGF-β superfamily in metabolic disease.

**Figure S1. Activin pathway components are dysregulated in mouse models of metabolic disease. (A-C)** mRNA expression by single nucleus RNA-seq (*28*) collected from C57Bl/6J mice on a standard chow diet. **(A)** Expression of type I receptors (*Acvr1b*, *Tgfbr1*, *Acvr1c*) and type II receptors (*Acvr2a*, *Acvr2b*, *Bmpr2*) in adipocytes (*28*) **(B)** UMAP representation of *Acvr1c* across adipose tissue (*28*) **(C)** *Acvr1c* expression distribution across distinct adipocyte subpopulations (*28*) **(D)** Adipocyte *Acvr1c* expression in adipose tissue from low-fat diet and high-fat diet (HFD) mice. **(E)** TPM of *Gdf3* expression in adipose tissue from chow diet and HFD mice (*28*). **(F)** TPM of *Inhbe* expression in the liver from normal, NAFLD, and fatty liver mice (*86–95*). Data in (A–C) are shown as violin plots; data in (D) are shown as a violin plot; data in (F) are shown as a box-and-whisker plot (median, interquartile range, min to max). P values determined by unpaired two-tailed t test for two-group comparisons (D, E) and ordinary one-way ANOVA with Dunnett’s multiple comparisons test vs. the normal control group (F).

**Figure S2. Activin E and GDF3 regulate lipolysis through ALK7-dependent SMAD signaling. (A)** Adipocyte differentiation marker gene expression in immortalized human adipocytes treated with recombinant ActE (0, 75, 150, 300 ng/ml) for 24 hours. **(B)** Lipolysis marker gene expression in immortalized human adipocytes treated with recombinant activin E (0, 75, 150, 300 ng/ml) for 24 hours**. (C&D)** Isoproterenol-stimulated free fatty acid release from immortalized mouse adipocytes pretreated with ALK5 inhibitors Reposox and Galunisertib (2.5, 5, 10 nM) for 4 hours prior to treatment with recombinant GDF3 (500 ng/ml) for 24 hours, normalized to controls. **(E)** mRNA levels in mouse immortalized adipocytes transduced with a short hairpin to a scrambled sequence (Scr) and overexpressing a constitutive *Gdf3* mRNA (*Gdf3*-OE). **(F)** TPM-normalized expression of type I and type II activin receptors expressed in HEK 293 cells (*96*). **(G)** Relative mean fluorescence intensity (MFI) of SBE-GFP in HEK293 reporter cells transiently transfected with indicated plasmids (n = 3 biological replicates per condition; each n represents average relative MFI of approximately 2000 sorted live cells). Reporter cells expressing plasmid-encoded mouse *Gdf3* and/or wild-type *ALK7* or SB431542-resistant *ALK4-ST*, *ALK5-ST*, or *ALK7-ST*, treated with or without SB431542. Data are mean ± SEM. P values for two-group comparisons (E) were determined by unpaired two-tailed t test. P values for comparisons of three or more groups (A, B, C, D) were determined by ordinary one-way ANOVA with Dunnett’s multiple comparisons test vs. the control group.

**Figure S3. eWAT expression profiling.**Bulk RNA-seq expression profiling of eWAT from mice on HFD from *Gdf3^fl/fl^*, *Gdf3^KO^*, *Gdf3^fl/fl^*:*Inhbe*^KD^, and *Gdf3^KO^*::*Inhbe*^KD^. **(A)** Genes involved in transcriptional regulation or enzymatic control of lipolysis. **(B)** Differentially expressed genes (DEG) between the four groups.

**Figure S4. *Gdf3* deletion with BIMA suppresses hepatic fibrosis and gluconeogenic gene expression. (A)** Quantification of liver fibrotic area (Sirius red staining) in *Gdf3*^fl/fl^ and *Gdf3*^KO^ mice with or without hepatocyte *Inhbe* knockdown (*Inhbe*^KD^). **(B)** *Gdf3* mRNA expression in adipose tissue from *Gdf3*f^l/fl-BIMA^ and *Gdf3*^KO-BIMA^ mice. **(C)** Expression of fibrosis marker genes in liver tissue from *Gdf3*^fl/fl-BIMA^ and *Gdf3*^KO-BIMA^ mice. **(D)** Expression of gluconeogenesis marker genes in liver tissue from *Gdf3*^fl/fl-BIMA^ and *Gdf3*^KO-BIMA^ mice. Data are mean ± SEM. P values in **(A)** were determined by ordinary two-way ANOVA (*Gdf3* genotype × *Inhbe*^KD^) with Tukey’s multiple comparisons test. P values in (B–D) were determined by an unpaired two-tailed Student’s t-test. *P < 0.05; **P < 0.01; ***P < 0.001; ****P < 0.0001; ns, not significant.

**Figure S5. *Gdf3* deficiency with BIMA attenuates hepatic fibrosis and gluconeogenesis in the thermoneutral Western diet model. (A)** mRNA expression of *Gdf3* in mouse adipose tissue. **(B)** Isoproterenol-stimulated free fatty acid release from immortalized mouse adipocytes treated with BIMA (15– 1000 µg/ml) for 24 hours, fold induction over untreated. (**C)** Isoproterenol-stimulated free fatty acid release from primary human adipose tissue slices from a patient with obesity treated with BIMA (50, 250 µg/ml) for 24 hours. **(D)** Quantification of liver fibrotic area (Sirius red staining) in *Gdf3*^fl/fl^ and *Gdf3*^KO^ mice with or without BIMA treatment, housed at thermoneutrality and fed a Western diet. **(E)** Expression of fibrosis marker genes in liver tissue from *Gdf3^fl/fl^*^-BIMA^ and *Gdf3^KO^*^-BIMA^ mice housed at thermoneutrality and fed a Western diet. **(F)** Expression of extracellular matrix (collagen) marker genes in liver tissue from *Gdf3^fl/fl^*^-BIMA^ and *Gdf3^KO^*^-BIMA^ mice housed at thermoneutrality and fed a Western diet. (**G)** Expression of gluconeogenesis marker genes in liver tissue from *Gdf3^fl/fl^*^-BIMA^ and *Gdf3^KO^*^-BIMA^ mice housed at thermoneutrality and fed a Western diet. Data are mean ± SEM. P values in (A) and (D) were determined by ordinary two-way ANOVA (*Gdf3* genotype × BIMA treatment). P values in (B) were determined by ordinary one-way ANOVA with Dunnett’s multiple comparisons test vs. control. P values in (C) and (E–G) were determined by ordinary one-way ANOVA with Tukey’s multiple comparisons test. *P < 0.05; **P < 0.01; ***P < 0.001; ****P < 0.0001; ns, not significant.

## MATERIALS AND METHODS

### Animal models and experimental design

All animal experiments were conducted in accordance with protocols approved by the Institutional Animal Care and Use Committee (IACUC) of the Beth Israel Deaconess Medical Center. Mice were housed in individually ventilated cages under standard conditions: 12 h light/12 h dark cycles (6am/6pm), ambient temperature of 22 ± 2 °C (except where otherwise specified), 30–70% humidity, with ad libitum access to food and water unless stated otherwise. Cages and bedding were changed biweekly, and mice were monitored daily by animal care technicians with veterinary oversight. All mice remained free of adventitious infections throughout the study duration.

### Generation of *Gdf3* conditional knockout mice

Homozygous *Gdf3* floxed mice (*Gdf3*^fl/fl^) were generated as described (*15*). Whole-body inducible *Gdf3* knockout (*Gdf3*^KO^) mice were generated by crossing homozygous *Gdf3*^fl/fl^ mice with hemizygous *Rosa26-Cre^ERT2/-^*mice (The Jackson Laboratory, stock no. 008463). Age-matched homozygous *Gdf3*^fl/fl^ littermates lacking the *Rosa26-Cre^ERT2^* transgene served as controls for all experiments. Both male and female mice were used as indicated in individual experiments.

### Dietary interventions and tamoxifen-induced gene deletion

Eight-week-old *Gdf3^fl/fl^* and *Gdf3^fl/fl^:: Rosa26-Cre^ERT2^* mice were fed either a high-fat diet (HFD; 60% kcal from fat; Research Diets, no. D12492i) for 8 weeks, or HFD followed by a Western diet (WD; 40% kcal from fat, primarily palm oil; 20% kcal from fructose; 2% cholesterol; Research Diets no. D09100310i) for 8 weeks. To induce *Gdf3* deletion, mice received intraperitoneal (IP) injections of tamoxifen (75 mg/kg body weight in 100% corn oil) daily for seven consecutive days. The mice then underwent a 4-week washout period after the final tamoxifen injection before subsequent interventions. Both groups of mice, *Gdf3^fl/fl^* and *Gdf3^fl/fl^::Rosa26-Cre^ERT2^*, received tamoxifen.

### si*Inhbe* and BIMA interventions

*Inhbe^KD^* mice received subcutaneous injections of si*Inhbe* (Eli Lilly) at 3 mg/kg body weight, twice weekly for 2 weeks to induce Activin E knockdown. Activin receptor blockade with a murine-Fc fragment, BYM338 (20 mg/kg body weight, twice weekly for 2 or 4 weeks as indicated. Vehicle-treated control mice received equivalent volumes of PBS.

To model diet-induced metabolic dysfunction under thermoneutral conditions, mice were initially fed a high-fat diet, followed by tamoxifen-induced Gdf3 knockout. After a 4-week washout period, mice were maintained at 30°C on a Western diet for 8-10 weeks and treated with Bimagrumab (20 mg/kg) for 2 weeks.

### Body composition

Fat mass and lean mass were quantified using an EchoMRI-100 body composition analyzer according to the manufacturer’s instructions. Conscious mice were placed in a restraint tube and scanned for approximately 90 s per measurement.

### Glucose tolerance tests

Mice were fasted for 4 h before receiving an IP injection of D-glucose (Sigma-Aldrich) at 1 g/kg body weight. Blood glucose concentrations were measured from tail vein samples at 0, 15, 30, 60, and 120 min post-injection using a handheld glucometer (Contour Next, Ascensia Diabetes Care).

### Insulin tolerance tests

Mice were fasted for 4 h before receiving an IP injection of Novolin R (Novo Nordisk), short-acting human recombinant insulin at 0.75 U/kg body weight. Blood glucose concentrations were measured at 0, 15, 30, 60, and 120 min post-injection.

### Pyruvate tolerance tests

Mice were fasted for 4 h before receiving an IP injection of sodium pyruvate (Sigma-Aldrich) at 2.5 g/kg body weight. Blood glucose concentrations were measured at 0, 15, 30, 60, and 120 min post-injection to assess hepatic gluconeogenic capacity.

### Indirect calorimetry

Metabolic rate measurements were performed using the Promethion indirect calorimetry system (Sable Systems International) with temperature and light-controlled chambers. Individually housed mice were acclimated to the chambers for approximately 24 h before data collection. Mice had ad libitum access to food and water and were maintained under 12 h light/12 h dark cycles (06:00/18:00) at 23 ± 0.2 °C unless otherwise specified. Positional tracking and physical activity were recorded every second. Oxygen consumption (VO_₂_) and carbon dioxide production (VCO_₂_) were measured every 3 min. Energy expenditure (EE) was calculated using the Weir equation (*97*). Respiratory exchange ratio (RER) was calculated as VCO_₂_/VO_₂_. Data were exported using the Macro Interpreter and analyzed using CalR version 2 (*98*). Automatic data cleaning (the “Remove Outliers” function) was performed to remove gross outliers (±3 SD from group mean) from malfunctioning cages. Food caching events greater than 100 mg/min were also removed. Energy balance was calculated as energy intake minus total energy expenditure.

### In vivo lipolysis

Mice were fasted for 4 h before receiving an IP injection of isoproterenol hydrochloride (10 mg/kg body weight; Sigma-Aldrich). Blood samples were collected via lateral tail vein puncture into EDTA-coated tubes at 0, 15, 30, 45, and 60 min post-injection. Samples were immediately placed on ice, centrifuged at 2,000*g* for 15 min at 4 °C, and plasma was stored at −80 °C until analysis. Plasma non-esterified fatty acid (NEFA) concentrations were quantified using a colorimetric assay kit (Free Fatty Acid Quantitation Kit, Abcam, no. ab65341) according to the manufacturer’s instructions.

### Ex vivo lipolysis

Epididymal white adipose tissue (eWAT) and inguinal white adipose tissue (iWAT) depots were rapidly excised and weighed. Approximately 70 mg of tissue from each depot was transferred to individual wells of a 24-well plate containing 1 ml pre-warmed Krebs–Ringer bicarbonate HEPES (KRBH) buffer (pH 7.4, 37 °C). Tissue samples were minced into small fragments using sterile scissors, and the medium was replaced with 1 ml KRBH buffer supplemented with 2% fatty acid-free bovine serum albumin (KRBH-BSA). After equilibration, the medium was replaced with 1 ml KRBH-BSA containing isoproterenol (10 μM). Plates were incubated at 37 °C for 5 min, after which 100 μl of medium was collected as the baseline sample and replaced with 100 μl fresh KRBH-BSA containing isoproterenol (10 μM). Plates were incubated at 37 °C for 2 h, after which all medium was collected. Both baseline and 2 h samples were heat-inactivated at 65 °C for 10 min to eliminate residual enzymatic activity and stored at −80 °C until analysis. Tissue samples were washed with phosphate-buffered saline (PBS) and lysed in RIPA buffer for protein quantification using the Pierce BCA Protein Assay Kit (Thermo Fisher Scientific). Free fatty acid (FFA) concentrations in the medium were measured using a colorimetric assay kit (Abcam, no. ab65341) and normalized to total protein content.

### Serum insulin

Blood samples were collected from the lateral tail vein after a 4 h fast into EDTA-coated tubes, immediately placed on ice, and centrifuged at 2,000*g* for 15 min at 4 °C. Plasma was stored at −80 °C until analysis. Insulin concentrations were measured using an Ultra-Sensitive Mouse Insulin ELISA Kit (Crystal Chem) according to the manufacturer’s protocol.

### Serum liver enzymes

Plasma alanine aminotransferase (ALT), aspartate aminotransferase (AST), and gamma-glutamyl transferase (GGT) activities were measured using colorimetric assay kits (Teco Diagnostics) according to the manufacturer’s instructions.

### Hepatic triglycerides

Liver tissue (approximately 50 mg) was homogenized in 5% NP-40 solution using a bead homogenizer and centrifuged at 10,000*g* for 10 min at 4°C. Supernatants were collected, and triglyceride concentrations were determined using a Triglyceride Colorimetric Assay Kit (Cayman Chemical) according to the manufacturer’s protocol. Results were normalized to protein concentration.

### Hepatic glycogen

Liver tissue (approximately 50 mg) was homogenized in distilled water, boiled for 5 min, and centrifuged at 10,000*g* for 5 min at 4 °C. Supernatants were collected, and glycogen concentrations were measured using a Glycogen Assay Kit (Cayman Chemical) according to the manufacturer’s instructions. Results were normalized to protein concentration.

### Histology

Liver, eWAT, and iWAT were fixed in 4% paraformaldehyde overnight at 4 °C for histological processing and stained with hematoxylin and eosin (H&E) or Sirius Red for collagen visualization. Images were acquired at ×20 magnification using a SLIDEVIEW VS200 digital slide scanner (Olympus). Adipocyte area was quantified using the Adiposoft plugin for ImageJ software (National Institutes of Health). For each sample, at least 500 adipocytes from multiple fields were analyzed, and the mean adipocyte area was calculated.

### RNA isolation and quantitative real-time PCR

Total RNA was extracted from snap-frozen tissues using the Direct-zol RNA Miniprep Kit (Zymo Research, no. R2050) according to the manufacturer’s protocol, including on-column DNase I digestion. RNA concentration and purity were assessed using a NanoDrop spectrophotometer (Thermo Fisher Scientific). Complementary DNA (cDNA) was synthesized from 1 μg total RNA using the High-Capacity cDNA Reverse Transcription Kit (Thermo Fisher Scientific, no. 4368813) according to the manufacturer’s instructions.

Quantitative real-time PCR (RT-qPCR) was performed using SYBR Select Master Mix (Applied Biosystems, no. 4472920) on a QuantStudio 6 Real-Time PCR System (Applied Biosystems). Reactions were performed in technical duplicate or triplicate in 10 μl volumes containing 5 μl SYBR Select Master Mix, 0.5 μl each of forward and reverse primers (10 μM), 2 μl cDNA template (diluted 1:10), and 2 μl nuclease-free water. Thermal cycling conditions were: 50 °C for 2 min, 95 °C for 2 min, followed by 40 cycles of 95 °C for 15 s and 60 °C for 1 min. Melt curve analysis was performed to verify amplicon specificity. Relative mRNA expression was calculated using the 2[−ΔΔCt] method, with normalization to *Tbp* (TATA-box binding protein) as the endogenous control.

### Human transcriptomic and proteomic data analysis

Publicly available transcriptomic datasets were obtained as follows: human and mouse single nucleus RNA-seq of adipose tissue from Emont et al (*28*) was accessed through the Broad Single Cell Portal (*99*). Data was also accessed from the Gene Expression Omnibus (GEO) for the Metabolic Syndrome in Men (METSIM) project across Finnish individuals of varying BMI (*29*), GSE135134. Data from GSE141432 were accessed to visualize changes in bulk adipose tissue RNA-seq in obesity and T2D (*83*). Similarly bulk adipose tissue RNA-seq data in weight loss was accessed from GSE59034 (*84*). Liver bulk RNA-seq was accessed from GSE126848 (*35*). Untargeted serum proteomic data across liver disease progression were obtained from the ProteomeXchange Consortium (dataset identifier PXD051911) (*85*). Data visualization and statistical analyses were performed using R with appropriate Bioconductor packages.

### Cell Lines

HEK293T cells (human female in origin) and HEK293 cells stably expressing the gene for Green Fluorescent Protein (GFP) downstream of the CAGA_12_ promoter (SBE-GFP) were cultured every 3–4 days using 0.05% Trypsin EDTA (Gibco, # 25300054) and maintained in DMEM (Gibco, # 11965118) supplemented with 10% iFBS and 1X penstrep. Immortalized Human and mouse iWAT-SVF cells were a gift from Shingo Kajimura’s lab. These cells were cultured every 3-4 days using 0.25% Trypsin-EDTA, and maintained in DMEM containing 10% iFBS and 1X pen-strep. All cell lines were maintained under sterile conditions at 37°C and 5% CO_2_.

### Human and Mouse *In Vitro* Adipocyte Differentiation

Immortalized preadipocytes were plated in collagen-coated plates and grown to confluence in standard cell-culture-treated dishes for mice and collagen-coated plates for humans. Differentiation was initiated (day 1) with the complete media containing the induction cocktail of 0.5mM 3-isobutyl-1-methylxanthine (IBMX, Sigma-Aldrich, # I5879), 2µg/ml dexamethasone (Sigma-Aldrich, # D4902-100MG), 125µM indomethacin (Sigma-Aldrich, # I7378), and 0.5μM rosiglitazone (Sigma-Aldrich, # R2408) for 2 days. The cells were then switched to maintenance media containing 10 µg/ml (Human) and 5 µg/ml (Mouse) insulin (Sigma-Aldrich, #I6634) and 1 µM (Human) and 0.5 µM (Mouse) rosiglitazone, which were refreshed every other day until day 10 or 7days or until 90 to 100% of cells were differentiated.

### *In Vitro* Lipolysis Assays

Immortalized preadipocytes were differentiated into mature adipocytes in 48-well or 12-well plates as described above, serum-starved for 4 hours in DMEM containing 1X pen-step and 0.3% fatty acid-free BSA, and then incubated with recombinant proteins ActE and GDF3 in serum-free medium for 24 hours prior to initiating lipolysis. The media were then aspirated from all wells and replaced with KRBH-BSA buffer. Baseline samples were collected immediately from all the wells and replaced with KRBH-BSA buffer containing isoproterenol (10 µM). Cells were incubated at 37 °C for the indicated time points, after which the media were collected (stimulated samples). All samples were stored at -80 °C until further processing. To measure FFA levels, all samples were thawed at room temperature and incubated at 65 °C for 10 min to inactivate any residual enzymatic activity. FFA levels were quantified using the Free Fatty Acid Quantitation Kit (Abcam # ab65341) following the manufacturer’s instructions.

### Flow cytometry Reporter Assays

SBE-GFP expressing reporter cells were plated overnight in 12 well plates to reach 50-70% confluency, transfected with plasmids encoding indicated genes using Lipofectamine 3000 (Invitrogen). Cells were allowed to grow overnight and then split into a 96 well collagen coated cell culture plate to reach 90% confluency overnight. Cells were then serum starved with or without the indicated recombinant proteins and/ or the small molecule inhibitor SB-431542 (10μM). After 24 hours, cells were processed for flow cytometry as described previously (*15*). Briefly, 96 well plate media was discarded, and cells were washed with PBS. Cells were then dissociated with 0.05% Trypsin EDTA at room temperature for approximately 5min. Trypsin was neutralized with triple the volume of PBS containing 0.3% fatty acid free BSA (FACS buffer) containing 1:10000 LIVE/DEAD Fixable Far Red Dead Cell Stain (Thermo Fisher Scientific, # L34973), the plate was incubated for 15 minutes on ice and the plate was spun in a cold centrifuge with a swinging rotor at 500g for 10min. The supernatant was discarded, the cells were washed with FACS buffer and centrifuged again. The supernatant was removed, and each well was incubated with 200uL of FACS buffer. These cells were then analyzed using the CytoFLEX flow Cytometer (Beckman Coulter) and the CytExpert acquisition and analysis software. Briefly, single unstained cells and single fluorescent controls were used to set the gains and gates for each fluorescent channel. Cells were gated to analyze at least 500 GFP positive live single cells from each well. The mean fluorescence intensity (MFI) of GFP of the live cells from each well was used for subsequent analysis.

### Study population and type 2 diabetes phenotype definition

Analyses were conducted in the UK Biobank (UKB), a prospective population cohort of approximately 500,000 adults aged 40–69 years across the United Kingdom (*39, 100*). Type 2 diabetes (T2D) case-control status was defined using the Eastwood algorithm (*101*) applied to linked primary care (GP) records, which integrates diagnostic Read codes, medication prescriptions, and laboratory measurements to adjudicate T2D status. Participants lacking GP record linkage or with ambiguous diabetes status under the algorithm were excluded. For this analysis, we included a total of 41,631 cases of T2D and 258,878 non-diabetes controls, and defined cases of T2D and controls using an algorithm designed specifically for the UKB. Of these, 25,827 T2D cases had complete covariate data and passed genotype quality-control filters and were retained for the burden association analysis described below. The UKB has obtained ethical approval covering the present study from the National Research Ethics Committee (REC ref. no. 11/NW/0382) and the data were accessed through application no. 27892.

### Predicted loss-of-function variant annotation and carrier identification

Predicted loss-of-function (pLoF) variants within ACVR1C and INHBE were extracted from UKB whole-exome sequencing data using BGENIX (v1.1.7), querying per-chromosome BGEN files over gene-level genomic intervals defined against the GRCh38 reference. Variants with minor allele count ≥ 1 in the analytic sample were retained. Variants were functionally annotated using SnpEff (v5.1) (*102*) against the GRCh38.99 database. High-impact pLoF consequences including stop-gain, frameshift, splice-donor, and splice-acceptor were retained for burden analysis. Variant dosages were extracted from UKB WES BGEN files via the ukbrap framework on the UKB Research Analysis Platform (DNAnexus). Carrier status was assigned as a binary indicator per gene, set to 1 if an individual carried at least one non-reference dosage call across any qualifying pLoF variant within that gene, and 0 otherwise.

### Genetic association analysis

Gene-level burden association with T2D was tested using logistic regression. Three nested covariate models were evaluated for each gene: Model 1 (base) included age, sex, BMI, and the first ten principal components of ancestry to account for population stratification; Model 2 additionally adjusted for waist-hip ratio (WHR); and Model 3 additionally adjusted for reticulocyte percentage (UKB Field 30240). Odds ratios (OR) and 95% confidence intervals were estimated from the exponentiated burden regression coefficient. Power was computed using the genpwr package (*103*) under a dominant genetic model (α = 0.05).

### Human adipose tissue preparation

Mesenteric adipose tissue was obtained from a female patient in her 60s with a BMI 36, undergoing combined liver-kidney transplantation for MASH cirrhosis and renal failure at Beth Israel Deaconess Medical Center (BIDMC). The patient was enrolled under the End Stage Liver Disease Patient Registry, and sample collection was approved by the BIDMC Institutional Review Board. Mesenteric adipose tissue was transported from the operating room in University of Wisconsin (UW) solution for immediate processing. For preparation of precision-cut adipose tissue slices, 8-mm tissue cores were embedded in 4% agarose and sectioned into 500-µm slices using a vibrating microtome (VF-510-0Z, Precisionary Instruments, MA) in UW solution. The slices were transferred to 24-well plates containing Dulbecco’s Modified Eagle Medium (DMEM), flushed with oxygen for 2 min, and incubated at 37 °C in 95% O_2_ and 5% CO_2_ with continuous orbital shaking at 150 rpm for 2 hours. The medium was exchanged daily under the same incubation conditions.

### Bulk RNA Sequencing of Adipose and Liver Tissues

For RNA isolation, total RNA was extracted from snap-frozen tissues using the Direct-zol RNA Miniprep Kit (Zymo Research, no. R2050) according to the manufacturer’s protocol, including on-column DNase I digestion. RNA concentration and purity were assessed using a NanoDrop spectrophotometer (Thermo Fisher Scientific).

Purified RNA samples were submitted to Plasmidsaurus (https://plasmidsaurus.com) for bulk RNA sequencing as a commercial service. Library preparation, sequencing, and initial quality control were performed by Plasmidsaurus, which delivered raw FASTQ files, gene-level count tables, and sequencing quality control reports. Sequencing yielded approximately 20 million raw reads per sample. Following deduplication, approximately 10 million deduplicated reads were retained per sample for downstream analysis.

### Data analysis

For data analysis, raw gene-level count tables were imported from tab-separated value (TSV) files containing Ensembl gene identifiers, gene symbols, and replicate-level count columns. Principal component analysis (PCA) was used for quality control and identification of outlier replicates, which were excluded from downstream analysis if total read counts were below 1 million reads. Samples were grouped into biologically defined experimental conditions prior to differential expression analysis. Pairwise differential expression analysis was performed using the DESeq2 package in R with default median-of-ratios normalization and dispersion estimation. Wald statistics were used for hypothesis testing, and Benjamini–Hochberg multiple-testing correction was applied to generate adjusted false discovery rate (FDR) q-values for all pairwise comparisons. Differentially expressed genes (DEGs) were defined as genes with an adjusted FDR < 0.10. Heatmap generation and downstream pathway visualization were performed using custom Python scripts utilizing pandas, NumPy, Matplotlib, and Seaborn. Expression matrices were transformed into log2 counts-per-million (log2CPM) values prior to normalization and visualization. Heatmaps were generated using condition-averaged expression profiles and visualized using row-wise min–max scaling. Hierarchical clustering was performed using average-linkage clustering with correlation distance metrics to identify transcriptional relationships among genes and experimental groups. For pathway enrichment analysis, DEGs passing the adjusted FDR < 0.10 threshold were analyzed using KEGG pathway enrichment through the g:Profiler Python API interface. Enrichment testing was restricted to KEGG pathways, and pathways with enrichment p-values < 0.05 were considered significantly enriched.

### Production of Lentiviral Vectors

Lentiviral particles were produced by transient three-plasmid co-transfection of HEK293T cells, a derivative of human embryonic kidney cells stably expressing the SV40 Large T antigen. The transfection system comprised the second-generation packaging plasmid psPAX2, the vesicular stomatitis virus glycoprotein (VSV-G) envelope plasmid pMD2.G, and one of two transfer plasmids: pLV[Exp]-CMV-mCherry-mGdf3, encoding murine Gdf3 under cytomegalovirus (CMV) promoter control with an mCherry fluorescent reporter; or pLV[shRNA]-TagBFP2-U6>Scramble_shRNA, expressing a non-targeting scramble shRNA control with a TagBFP2 reporter.

HEK293T cells were seeded onto 10 cm tissue culture dishes and cultured to 80–90% confluence prior to transfection. Plasmid DNA was diluted in Opti-MEM reduced-serum medium (Thermo Fisher Scientific) and complexed with linear polyethylenimine (PEI; molecular weight 25,000 Da; Polysciences Inc.) at a 1:3 DNA: PEI mass ratio. Transfection complexes were incubated for 15 min at room temperature before dropwise addition to cells. Culture medium was replaced 16–18 h post-transfection with 10 mL fresh Dulbecco’s Modified Eagle Medium (DMEM) supplemented with 10% (v/v) fetal bovine serum (FBS). Viral supernatants were harvested at 48, 72, and 96 h post-transfection. Each harvest was centrifuged at 500 × g for 5 min at 4°C to pellet cellular debris, and the clarified supernatant was filtered through a 0.45 µm polyethersulfone (PES) membrane filter (MilliporeSigma). Filtered supernatants from all three time points were pooled, aliquoted, and stored at −80°C until use.

### Lentiviral Transduction of Adipocytes

Adipocyte differentiation was initiated on day 1 by replacing growth medium with complete adipogenic induction medium consisting of DMEM supplemented with 10% (v/v) FBS, 0.5 mM 3-isobutyl-1-methylxanthine (IBMX; Sigma-Aldrich, cat. no. I5879), 2 µg/mL dexamethasone (Sigma-Aldrich, cat. no. D4902), 125 µM indomethacin (Sigma-Aldrich, cat. no. I7378), and 0.5 µM rosiglitazone (Sigma-Aldrich, cat. no. R2408) . Cells were maintained in induction medium for 2 days (days 1–2).On day 3, cells were switched to maintenance medium and lentiviral transduction was initiated concurrently. For human adipocytes, maintenance medium consisted of DMEM supplemented with 10% FBS, 10 µg/mL recombinant human insulin (Sigma-Aldrich, cat. no. I6634), and 1 µM rosiglitazone. For mouse adipocytes, maintenance medium consisted of DMEM supplemented with 10% FBS, 5 µg/mL recombinant mouse insulin, and 0.5 µM rosiglitazone. Lentiviral supernatant was added directly to the respective maintenance medium in the presence of 5 µg/mL polybrene (hexadimethrine bromide; MilliporeSigma) to enhance transduction efficiency. The medium containing the virus and polybrene was replaced daily with freshly prepared maintenance medium supplemented with lentiviral supernatant and polybrene for 5 consecutive days (days 3–7).

### Statistical Analyses

Data analysis and plots were generated using GraphPad Prism software or the R programming language version 4.4.2. Indirect calorimetry data were exported with Macro Interpreter, macro 13 (Sable Systems), before analysis in CalR version 2 according to the internationally accepted standards (*48, 98*). All data are expressed as mean values ± SEM unless specified otherwise. All data were assumed to have a normal distribution, and two-tailed Student’s t-tests, one-way ANOVA with Fisher’s LSD Multiple Comparison tests, two-way or three-way ANOVA were used to compare means between groups; p < 0.05 was considered significant.

## Acknowledgements

This work is supported by NIH grants R01DK107717, RC2DK142612, P30DK135043 to ASB and an investigator-initiated research grant from Eli Lilly and Company to ASB. NIH grant T32DK007516 to support JAH. J.M.M. is supported by American Diabetes Association grant #11-22-ICTSPM-16 and by NHGRI U01HG011723, by the National Institute Of Diabetes And Digestive And Kidney Diseases of the National Institutes of Health under Award Number R01DK137993, R01DK140545 and U01 DK140757, AMP CMD award from RFP 6 from the Foundation for the National Institutes of Health, and a Medical University of Bialystok (MUB) grant from the Ministry of Science and Higher Education (Poland). This work is supported by the Novo Nordisk Foundation (NNF21SA0072102).

## Association of LoF variants in ACVR1C and INHBE with T2D in UKBB

pLoF variants were aggregated per gene and tested for association with T2D case/control status using logistic regression (25,827 cases, 258,878 controls). Three covariate models were tested on Model 1 (base): age, sex, BMI, and PC1–10; Model 2: Model 1 additionally adjusted for waist-hip ratio; Model 3: Model 1 additionally adjusted for reticulocyte percentage. Variants (n): number of LoF variants included in the burden. Total carriers: number of individuals carrying ≥1 LoF allele. Median and maximum carriers per variant describe the carrier count distribution across aggregated variants. OR and 95% CI were derived from the burden regression coefficient. Power was estimated at the observed OR and carrier frequency for each model using a dominant genetic model at α = 0.05. Power (OR=0.15) represents power at a hypothetical stronger effect size for ACVR1C; dash (–) indicates this column was not applicable for INHBE given adequate observed power., pLoF predicted loss of function

---

**Supplemental Table 1.** Association of LoF variants in ACVR1C and INHBE with T2D in UKBB. pLoF variants were aggregated per gene and tested for association with T2D case/control status using logistic regression (25,827 cases, 258,878 controls). Three covariate models were tested on Model 1 (base): age, sex, BMI, and PC1–10; Model 2: Model 1 additionally adjusted for waist-hip ratio; Model 3: Model 1 additionally adjusted for reticulocyte percentage. Variants (n): number of LoF variants included in the burden. Total carriers: number of individuals carrying ≥1 LoF allele. Median and maximum carriers per variant describe the carrier count distribution across aggregated variants. OR and 95% CI were derived from the burden regression coefficient. Power was estimated at the observed OR and carrier frequency for each model using a dominant genetic model at α = 0.05. Power (OR=0.15) represents power at a hypothetical stronger effect size for ACVR1C; dash (–) indicates this column was not applicable for INHBE given adequate observed power., pLoF predicted loss of function

| Gene | variants (n) | Total carriers | Median carriers | Maximum carriers | Model | OR (95% CI) | P value | Power | Power (OR=0.15) |
| --- | --- | --- | --- | --- | --- | --- | --- | --- | --- |
| ACVR1C | 21 | 38 | 1 | 12 | Model 1<br>(age, sex, BMI, PCs) | 0.307<br>(0.040–2.332) | 0.254 | 58.60% | 80.50% |
|  |  |  |  |  | Model 2<br>(+WHR) | 0.292<br>(0.038–2.278) | 0.24 | 60.90% | 80.40% |
|  |  |  |  |  | Model 3<br>(+Reticulocyte %) | 0.305<br>(0.040–2.328) | 0.252 | 57.00% | 78.70% |
| INHBE | 44 | 1,017 | 2-3 | 655 | Model 1<br>(age, sex, BMI, PCs) | 0.714<br>(0.544–0.935) | 0.015 | 98.10% | - |
|  |  |  |  |  | Model 2<br>(+WHR) | 0.767<br>(0.582–1.012) | 0.06 | 90.10% | - |
|  |  |  |  |  | Model 3<br>(+Reticulocyte %) | 0.736<br>(0.559–0.969) | 0.029 | 95.20% | - |

## Notes

### Competing Interest Statement

Stephen E. Flaherty 3rd, Paul M. Titchenell, Henning F. Kramer are employees of Eli Lilly. Alexander S. Banks is the recipient of an investigator initiated grant from Eli Lilly

## References

1. E. Fabbrini, S. Sullivan, S. Klein, Obesity and nonalcoholic fatty liver disease: biochemical, metabolic, and clinical implications. Hepatology 51, 679–89 (2010).

2. M.-J. Lee, Transforming growth factor beta superfamily regulation of adipose tissue biology in obesity. Biochim. Biophys. Acta Mol. Basis Dis. 1864, 1160–1171 (2018).

3. S. Modica, L. G. Straub, M. Balaz, W. Sun, L. Varga, P. Stefanicka, M. Profant, E. Simon, H. Neubauer, B. Ukropcova, J. Ukropec, C. Wolfrum, Bmp4 Promotes a Brown to White-like Adipocyte Shift. Cell Rep. 16, 2243–2258 (2016).

4. B. Gu, J. M. Linton, B. G. Hendrickson, H. Li, R. Hadas, G. Manella, J. Gregrowicz, B. Anggito, R. Lu, M. B. Elowitz, Programmable pathway profiles reveal signaling principles of TGF-β superfamily receptors (2026), doi:10.1101/2025.06.15.659618.

5. A. M. Deaton, A. Dubey, L. D. Ward, P. Dornbos, J. Flannick, E. Yee, S. Ticau, L. Noetzli, M. M. Parker, R. A. Hoffing, C. Willis, M. E. Plekan, A. M. Holleman, G. Hinkle, K. Fitzgerald, A. K. Vaishnaw, P. Nioi, Rare loss of function variants in the hepatokine gene INHBE protect from abdominal obesity. Nat. Commun. 13, 4319 (2022).

6. P. Akbari, O. A. Sosina, J. Bovijn, K. Landheer, J. B. Nielsen, M. Kim, S. Aykul, T. De, M. E. Haas, G. Hindy, N. Lin, I. R. Dinsmore, J. Z. Luo, S. Hectors, B. Geraghty, M. Germino, L. Panagis, P. Parasoglou, J. R. Walls, G. Halasz, G. S. Atwal, M. Jones, M. G. LeBlanc, C. D. Still, D. J. Carey, A. Giontella, M. Orho-Melander, J. Berumen, P. Kuri-Morales, J. Alegre-Díaz, J. M. Torres, J. R. Emberson, R. Collins, D. J. Rader, B. Zambrowicz, A. J. Murphy, S. Balasubramanian, J. D. Overton, J. G. Reid, A. R. Shuldiner, M. Cantor, G. R. Abecasis, M. A. R. Ferreira, M. W. Sleeman, V. Gusarova, J. Altarejos, C. Harris, A. N. Economides, V. Idone, K. Karalis, G. Della Gatta, T. Mirshahi, G. D. Yancopoulos, O. Melander, J. Marchini, R. Tapia-Conyer, A. E. Locke, A. Baras, N. Verweij, L. A. Lotta, Multiancestry exome sequencing reveals INHBE mutations associated with favorable fat distribution and protection from diabetes. Nat. Commun. 13, 4844 (2022).

7. C. A. Emdin, A. V. Khera, K. Aragam, M. Haas, M. Chaffin, D. Klarin, P. Natarajan, A. Bick, S. M. Zekavat, A. Nomura, DNA sequence variation in ACVR1C encoding the activin receptor-like kinase 7 influences body fat distribution and protects against type 2 diabetes. Diabetes 68, 226–234 (2019).

8. M. Koprulu, Y. Zhao, E. Wheeler, L. Dong, N. Rocha, C. Li, J. D. Griffin, S. Patel, M. Van de Streek, C. A. Glastonbury, I. D. Stewart, F. R. Day, J. Luan, N. Bowker, L. B. L. Wittemans, N. D. Kerrison, L. Cai, D. M. E. Lucarelli, I. Barroso, M. I. McCarthy, R. A. Scott, V. Saudek, K. S. Small, N. J. Wareham, R. K. Semple, J. R. B. Perry, S. O’Rahilly, L. A. Lotta, C. Langenberg, D. B. Savage, Identification of Rare Loss-of-Function Genetic Variation Regulating Body Fat Distribution. J. Clin. Endocrinol. Metab. 107, 1065–1077 (2022).

9. S. Y. Park, Y. Cho, S. M. Son, J. H. Hur, Y. Kim, H. Oh, H. Y. Lee, S. Jung, S. Park, I. Y. Kim, S. J. Lee, C. S. Choi, Activin E is a new guardian protecting against hepatic steatosis via inhibiting lipolysis in white adipose tissue. Exp Mol Med 57, 466–477 (2025).

10. R. C. Adam, D. S. Pryce, J. S. Lee, Y. Zhao, I. J. Mintah, S. Min, G. Halasz, J. Mastaitis, G. S. Atwal, S. Aykul, V. Idone, A. N. Economides, L. A. Lotta, A. J. Murphy, G. D. Yancopoulos, M. W. Sleeman, V. Gusarova, Activin E–ACVR1C cross talk controls energy storage via suppression of adipose lipolysis in mice. Proc. Natl. Acad. Sci. 120, e2309967120 (2023).

11. S. Yogosawa, S. Mizutani, Y. Ogawa, T. Izumi, Activin receptor-like kinase 7 suppresses lipolysis to accumulate fat in obesity through downregulation of peroxisome proliferator-activated receptor gamma and C/EBPalpha. Diabetes 62, 115–23 (2013).

12. J. D. Griffin, J. M. Buxton, J. A. Culver, R. Barnes, E. A. Jordan, A. R. White, S. E. Flaherty, B. Bernardo, T. Ross, K. K. Bence, M. J. Birnbaum, Hepatic Activin E mediates liver-adipose inter-organ communication, suppressing adipose lipolysis in response to elevated serum fatty acids. Mol Metab 78, 101830 (2023).

13. M. Sakaki, T. Shikata, K. Aoki, M. Takano, A. Kurisaki, M. Funaba, O. Hashimoto, Involvement of Activin E depletion in metabolic dysfunction-associated steatohepatitis. Biochem. Biophys. Rep. 44, 102339 (2025).

14. P. Tangseefa, H. Jin, H. Zhang, M. Xie, C. F. Ibáñez, Human ACVR1C missense variants that correlate with altered body fat distribution produce metabolic alterations of graded severity in knock-in mutant mice. Mol Metab 81, 101890 (2024).

15. N. Kotikalapudi, D. Ramachandran, D. Vieira, W. B. Rubio, G. R. Gipson, L. Troncone, K. Vestal, D. E. Maridas, V. Rosen, P. B. Yu, T. B. Thompson, A. S. Banks, Acute regulation of murine adipose tissue lipolysis and insulin resistance by the TGFβ superfamily protein GDF3. Nat. Commun. 16, 4432 (2025).

16. A. J. Levine, Z. J. Levine, A. H. Brivanlou, GDF3 is a BMP inhibitor that can activate Nodal signaling only at very high doses. Dev. Biol. 325, 43–48 (2009).

17. A. J. Levine, A. H. Brivanlou, GDF3, a BMP inhibitor, regulates cell fate in stem cells and early embryos. Dev. Camb. Engl. 133, 209–216 (2006).

18. O. Andersson, P. Bertolino, C. F. Ibáñez, Distinct and cooperative roles of mammalian Vg1 homologs GDF1 and GDF3 during early embryonic development. Dev Biol 311, 500–511 (2007).

19. O. Andersson, M. Korach-Andre, E. Reissmann, C. F. Ibanez, P. Bertolino, Growth/differentiation factor 3 signals through ALK7 and regulates accumulation of adipose tissue and diet-induced obesity. Proc Natl Acad Sci U A 105, 7252–6 (2008).

20. Y. Bu, K. Okunishi, S. Yogosawa, K. Mizuno, M. J. Irudayam, C. W. Brown, T. Izumi, Insulin Regulates Lipolysis and Fat Mass by Upregulating Growth/Differentiation Factor 3 in Adipose Tissue Macrophages. Diabetes 67, 1761–1772 (2018).

21. C. Chen, S. M. Ware, A. Sato, D. E. Houston-Hawkins, R. Habas, M. M. Matzuk, M. M. Shen, C. W. Brown, The Vg1-related protein Gdf3 acts in a Nodal signaling pathway in the pre-gastrulation mouse embryo. Development 133, 319–329 (2006).

22. J. A. Hall, D. Ramachandran, H. C. Roh, J. R. DiSpirito, T. Belchior, P. J. H. Zushin, C. Palmer, S. Y. Hong, A. I. Mina, B. Y. Liu, Z. M. Deng, P. Aryal, C. Jacobs, D. Tenen, C. W. Brown, J. F. Charles, G. I. Shulman, B. B. Kahn, L. T. Y. Tsai, E. D. Rosen, B. M. Spiegelman, A. S. Banks, Obesity-Linked PPARγ S273 Phosphorylation Promotes Insulin Resistance through Growth Differentiation Factor 3. Cell Metab 32, 665-+ (2020).

23. J. H. Choi, A. S. Banks, J. L. Estall, S. Kajimura, P. Bostrom, D. Laznik, J. L. Ruas, M. J. Chalmers, T. M. Kamenecka, M. Bluher, P. R. Griffin, B. M. Spiegelman, Anti-diabetic drugs inhibit obesity-linked phosphorylation of PPARgamma by Cdk5. Nature 466, 451–6 (2010).

24. S. J. Choi, Z. Yablonka-Reuveni, K. J. Kaiyala, K. Ogimoto, M. W. Schwartz, B. E. Wisse, Increased energy expenditure and leptin sensitivity account for low fat mass in myostatin-deficient mice. Am. J. Physiol.-Endocrinol. Metab. 300, E1031–E1037 (2011).

25. A. S. Banks, F. E. McAllister, J. P. Camporez, P. J. Zushin, M. J. Jurczak, D. Laznik-Bogoslavski, G. I. Shulman, S. P. Gygi, B. M. Spiegelman, An ERK/Cdk5 axis controls the diabetogenic actions of PPARgamma. Nature 517, 391–5 (2015).

26. J. Zhong, D. Zareifi, S. Weinbrenner, M. Hansen, F. Klingelhuber, P. A. Nono Nankam, S. Frendo-Cumbo, N. Bhalla, L. Cordeddu, T. de Castro Barbosa, P. Arner, I. Dahlman, M. Muniandy, S. Heinonen, K. H. Pietiläinen, A. Hoffmann, A. Ghosh, D. John, A. Tönjes, P. L. Ståhl, Y. Böttcher, M. Keller, P. Kovacs, A. G. Kerr, D. Langin, C. Wolfrum, M. Blüher, N. Krahmer, L. Massier, N. Mejhert, M. Rydén, adiposetissue.org: A knowledge portal integrating clinical and experimental data from human adipose tissue. Cell Metab. 37, 566– 569 (2025).

27. M. Civelek, Y. Wu, C. Pan, C. K. Raulerson, A. Ko, A. He, C. Tilford, N. K. Saleem, A. Stančáková, L. J. Scott, C. Fuchsberger, H. M. Stringham, A. U. Jackson, N. Narisu, P. S. Chines, K. S. Small, J. Kuusisto, B. W. Parks, P. Pajukanta, T. Kirchgessner, F. S. Collins, P. S. Gargalovic, M. Boehnke, M. Laakso, K. L. Mohlke, A. J. Lusis, Genetic Regulation of Adipose Gene Expression and Cardio-Metabolic Traits. Am. J. Hum. Genet. 100, 428–443 (2017).

28. M. P. Emont, C. Jacobs, A. L. Essene, D. Pant, D. Tenen, G. Colleluori, A. Di Vincenzo, A. M. Jørgensen, H. Dashti, A. Stefek, E. McGonagle, S. Strobel, S. Laber, S. Agrawal, G. P. Westcott, A. Kar, M. L. Veregge, A. Gulko, H. Srinivasan, Z. Kramer, E. De Filippis, E. Merkel, J. Ducie, C. G. Boyd, W. Gourash, A. Courcoulas, S. J. Lin, B. T. Lee, D. Morris, A. Tobias, A. V. Khera, M. Claussnitzer, T. H. Pers, A. Giordano, O. Ashenberg, A. Regev, L. T. Tsai, E. D. Rosen, A single-cell atlas of human and mouse white adipose tissue. Nature 603, 926–933 (2022).

29. M. Laakso, J. Kuusisto, A. Stančáková, T. Kuulasmaa, P. Pajukanta, A. J. Lusis, F. S. Collins, K. L. Mohlke, M. Boehnke, The Metabolic Syndrome in Men study: a resource for studies of metabolic and cardiovascular diseases. J. Lipid Res. 58, 481–493 (2017).

30. L. M. Carlsson, P. Jacobson, A. Walley, P. Froguel, L. Sjöström, P. A. Svensson, K. Sjöholm, ALK7 expression is specific for adipose tissue, reduced in obesity and correlates to factors implicated in metabolic disease. Biochem Biophys Res Commun 382, 309–14 (2009).

31. C. A. Emdin, A. V. Khera, K. Aragam, M. Haas, M. Chaffin, D. Klarin, P. Natarajan, A. Bick, S. M. Zekavat, A. Nomura, D. Ardissino, J. G. Wilson, H. Schunkert, R. McPherson, H. Watkins, R. Elosua, M. J. Bown, N. J. Samani, U. Baber, J. Erdmann, N. Gupta, J. Danesh, D. Saleheen, S. Gabriel, S. Kathiresan, DNA Sequence Variation in ACVR1C Encoding the Activin Receptor-Like Kinase 7 Influences Body Fat Distribution and Protects Against Type 2 Diabetes. Diabetes 68, 226–234 (2019).

32. A. E. Justice, T. Karaderi, H. M. Highland, K. L. Young, M. Graff, Y. Lu, V. Turcot, P. L. Auer, R. S. Fine, X. Guo, C. Schurmann, A. Lempradl, E. Marouli, A. Mahajan, T. W. Winkler, A. E. Locke, C. Medina-Gomez, T. Esko, S. Vedantam, A. Giri, K. S. Lo, T. Alfred, P. Mudgal, M. C. Y. Ng, N. L. Heard-Costa, M. F. Feitosa, A. K. Manning, S. M. Willems, S. Sivapalaratnam, G. Abecasis, D. S. Alam, M. Allison, P. Amouyel, Z. Arzumanyan, B. Balkau, L. Bastarache, S. Bergmann, L. F. Bielak, M. Blüher, M. Boehnke, H. Boeing, E. Boerwinkle, C. A. Böger, J. Bork-Jensen, E. P. Bottinger, D. W. Bowden, I. Brandslund, L. Broer, A. A. Burt, A. S. Butterworth, M. J. Caulfield, G. Cesana, J. C. Chambers, D. I. Chasman, Y.-D. I. Chen, R. Chowdhury, C. Christensen, A. Y. Chu, F. S. Collins, J. P. Cook, A. J. Cox, D. S. Crosslin, J. Danesh, P. I. W. de Bakker, S. de Denus, R. de Mutsert, G. Dedoussis, E. W. Demerath, J. G. Dennis, J. C. Denny, E. Di Angelantonio, M. Dörr, F. Drenos, M.-P. Dubé, A. M. Dunning, D. F. Easton, P. Elliott, E. Evangelou, A.-E. Farmaki, S. Feng, E. Ferrannini, J. Ferrieres, J. C. Florez, M. Fornage, C. S. Fox, P. W. Franks, N. Friedrich, W. Gan, I. Gandin, P. Gasparini, V. Giedraitis, G. Girotto, M. Gorski, H. Grallert, N. Grarup, M. L. Grove, S. Gustafsson, J. Haessler, T. Hansen, A. T. Hattersley, C. Hayward, I. M. Heid, O. L. Holmen, G. K. Hovingh, J. M. M. Howson, Y. Hu, Y.-J. Hung, K. Hveem, M. A. Ikram, E. Ingelsson, A. U. Jackson, G. P. Jarvik, Y. Jia, T. Jørgensen, P. Jousilahti, J. M. Justesen, B. Kahali, M. Karaleftheri, S. L. R. Kardia, F. Karpe, F. Kee, H. Kitajima, P. Komulainen, J. S. Kooner, P. Kovacs, B. K. Krämer, K. Kuulasmaa, J. Kuusisto, M. Laakso, T. A. Lakka, D. Lamparter, L. A. Lange, C. Langenberg, E. B. Larson, N. R. Lee, W.-J. Lee, T. Lehtimäki, C. E. Lewis, H. Li, J. Li, R. Li-Gao, L.-A. Lin, X. Lin, L. Lind, J. Lindström, A. Linneberg, C.-T. Liu, D. J. Liu, J. Luan, L.-P. Lyytikäinen, S. MacGregor, R. Mägi, S. Männistö, G. Marenne, J. Marten, N. G. D. Masca, M. I. McCarthy, K. Meidtner, E. Mihailov, L. Moilanen, M. Moitry, D. O. Mook-Kanamori, A. Morgan, A. P. Morris, M. Müller-Nurasyid, P. B. Munroe, N. Narisu, C. P. Nelson, M. Neville, I. Ntalla, J. R. O’Connell, K. R. Owen, O. Pedersen, G. M. Peloso, C. E. Pennell, M. Perola, J. A. Perry, J. R. B. Perry, T. H. Pers, A. Ewing, O. Polasek, O. T. Raitakari, A. Rasheed, C. K. Raulerson, R. Rauramaa, D. F. Reilly, A. P. Reiner, P. M. Ridker, M. A. Rivas, N. R. Robertson, A. Robino, I. Rudan, K. S. Ruth, D. Saleheen, V. Salomaa, N. J. Samani, P. J. Schreiner, M. B. Schulze, R. A. Scott, M. Segura-Lepe, X. Sim, A. J. Slater, K. S. Small, B. H. Smith, J. A. Smith, L. Southam, T. D. Spector, E. K. Speliotes, K. Stefansson, V. Steinthorsdottir, K. E. Stirrups, K. Strauch, H. M. Stringham, M. Stumvoll, L. Sun, P. Surendran, K. M. A. Swart, J.-C. Tardif, K. D. Taylor, A. Teumer, D. J. Thompson, G. Thorleifsson, U. Thorsteinsdottir, B. H. Thuesen, A. Tönjes, M. Torres, E. Tsafantakis, J. Tuomilehto, A. G. Uitterlinden, M. Uusitupa, C. M. van Duijn, M. Vanhala, R. Varma, S. H. Vermeulen, H. Vestergaard, V. Vitart, T. F. Vogt, D. Vuckovic, L. E. Wagenknecht, M. Walker, L. Wallentin, F. Wang, C. A. Wang, S. Wang, N. J. Wareham, H. R. Warren, D. M. Waterworth, J. Wessel, H. D. White, C. J. Willer, J. G. Wilson, A. R. Wood, Y. Wu, H. Yaghootkar, J. Yao, L. M. Yerges-Armstrong, R. Young, E. Zeggini, X. Zhan, W. Zhang, J. H. Zhao, W. Zhao, H. Zheng, W. Zhou, M. C. Zillikens, F. Rivadeneira, I. B. Borecki, J. A. Pospisilik, P. Deloukas, T. M. Frayling, G. Lettre, K. L. Mohlke, J. I. Rotter, Z. Kutalik, J. N. Hirschhorn, L. A. Cupples, R. J. F. Loos, K. E. North, C. M. Lindgren, CHD Exome+ Consortium, Cohorts for Heart and Aging Research in Genomic Epidemiology (CHARGE) Consortium, EPIC-CVD Consortium, ExomeBP Consortium, Global Lipids Genetic Consortium, GoT2D Genes Consortium, InterAct, ReproGen Consortium, T2D-Genes Consortium, MAGIC Investigators, Protein-coding variants implicate novel genes related to lipid homeostasis contributing to body-fat distribution. Nat. Genet. 51, 452–469 (2019).

33. M. Koprulu, Y. Zhao, E. Wheeler, L. Dong, N. Rocha, C. Li, J. D. Griffin, S. Patel, M. Van de Streek, C. A. Glastonbury, I. D. Stewart, F. R. Day, J. Luan, N. Bowker, L. B. L. Wittemans, N. D. Kerrison, L. Cai, D. M. E. Lucarelli, I. Barroso, M. I. McCarthy, R. A. Scott, V. Saudek, K. S. Small, N. J. Wareham, R. K. Semple, J. R. B. Perry, S. O’Rahilly, L. A. Lotta, C. Langenberg, D. B. Savage, Identification of Rare Loss-of-Function Genetic Variation Regulating Body Fat Distribution. J. Clin. Endocrinol. Metab. 107, 1065–1077 (2022).

34. M. Zhao, K. Okunishi, Y. Bu, O. Kikuchi, H. Wang, T. Kitamura, T. Izumi, Targeting activin receptor–like kinase 7 ameliorates adiposity and associated metabolic disorders. JCI Insight 8, e161229.

35. M. P. Suppli, K. T. G. Rigbolt, S. S. Veidal, S. Heebøll, P. L. Eriksen, M. Demant, J. I. Bagger, J. C. Nielsen, D. Oró, S. W. Thrane, A. Lund, C. Strandberg, M. J. Kønig, T. Vilsbøll, N. Vrang, K. L. Thomsen, H. Grønbæk, J. Jelsing, H. H. Hansen, F. K. Knop, Hepatic transcriptome signatures in patients with varying degrees of nonalcoholic fatty liver disease compared with healthy normal-weight individuals. Am J Physiol Gastrointest Liver Physiol 316, G462–G472 (2019).

36. G. Alvarez-Sola, N. Shrestha, R. Benedé-Ubieto, B. Jin, T. J. Kendall, J. A. Fallowfield, R. P. Goodman, LiRNA: An interactive atlas of human liver RNAseq databases. JHEP Rep. Innov. Hepatol. 8, 101914 (2026).

37. T. J. Kendall, M. Jimenez-Ramos, F. Turner, P. Ramachandran, J. Minnier, M. D. McColgan, M. Alam, H. Ellis, D. R. Dunbar, G. Kohnen, P. Konanahalli, K. A. Oien, L. Bandiera, F. Menolascina, A. Juncker-Jensen, D. Alexander, C. Mayor, I. N. Guha, J. A. Fallowfield, An integrated gene-to-outcome multimodal database for metabolic dysfunction-associated steatotic liver disease. Nat. Med. 29, 2939–2953 (2023).

38. Z. Li, H. Zhang, Q. Li, W. Feng, X. Jia, R. Zhou, Y. Huang, Y. Li, Z. Hu, X. Hu, X. Zhu, S. Huang, GepLiver: an integrative liver expression atlas spanning developmental stages and liver disease phases. Sci. Data 10, 376 (2023).

39. C. Bycroft, C. Freeman, D. Petkova, G. Band, L. T. Elliott, K. Sharp, A. Motyer, D. Vukcevic, O. Delaneau, J. O’Connell, A. Cortes, S. Welsh, A. Young, M. Effingham, G. McVean, S. Leslie, N. Allen, P. Donnelly, J. Marchini, The UK Biobank resource with deep phenotyping and genomic data. Nature 562, 203–209 (2018).

40. UK Biobank Whole-Genome Sequencing Consortium, Whole-genome sequencing of 490,640 UK Biobank participants. Nature 645, 692–701 (2025).

41. G. Manzi, E. Maggio, T. Recchioni, F. I. Adamo, A. Caputo, A. Mihai, S. Papa, G. Serino, G. Scoccia, R. Badagliacca, C. D. Vizza, The pleiotropic effects of sotatercept. Vascul. Pharmacol. 162, 107577 (2026).

42. E. English, I. Idris, G. Smith, K. Dhatariya, E. S. Kilpatrick, W. G. John, The effect of anaemia and abnormalities of erythrocyte indices on HbA1c analysis: a systematic review. Diabetologia 58, 1409–1421 (2015).

43. L. Ongaro, D. J. Bernard, Activin Actions in Adipocytes. J. Clin. Endocrinol. Metab. 110, 1803–1810 (2025).

44. L. J. Jonk, S. Itoh, C. H. Heldin, P. ten Dijke, W. Kruijer, Identification and functional characterization of a Smad binding element (SBE) in the JunB promoter that acts as a transforming growth factor-beta, activin, and bone morphogenetic protein-inducible enhancer. J Biol Chem 273, 21145–52 (1998).

45. K. A. Vestal, C. Kattamuri, M. Koyiloth, L. Ongaro, J. A. Howard, A. M. Deaton, S. Ticau, A. Dubey, D. J. Bernard, T. B. Thompson, Activin E is a transforming growth factor β ligand that signals specifically through activin receptor-like kinase 7. Biochem J 481, 547–564 (2024).

46. E. J. Goebel, L. Ongaro, E. C. Kappes, K. Vestal, E. Belcheva, R. Castonguay, R. Kumar, D. J. Bernard, T. B. Thompson, R. J. Davis, M. E. Bronner, A. P. Hinck, Eds. The orphan ligand, activin C, signals through activin receptor-like kinase 7. Elife 11, e78197 (2022).

47. E. J. Goebel, R. A. Corpina, C. S. Hinck, M. Czepnik, R. Castonguay, R. Grenha, A. Boisvert, G. Miklossy, P. T. Fullerton, M. M. Matzuk, V. J. Idone, A. N. Economides, R. Kumar, A. P. Hinck, T. B. Thompson, Structural characterization of an activin class ternary receptor complex reveals a third paradigm for receptor specificity. Proc. Natl. Acad. Sci. U. S. A. 116, 15505–15513 (2019).

48. A. S. Banks, D. B. Allison, T. Alquier, null Ansarullah, S. N. Austad, J. Auwerx, J. E. Ayala, J. A. Baur, S. Carobbio, G. A. Churchill, M. Dall, R. de Cabo, J. Donato, N. R. V. Dragano, C. F. Elias, A. W. Ferrante, B. N. Finck, J. E. Galgani, Z. Gerhart-Hines, L. J. Goodyear, J. L. Grobe, R. K. Gupta, K. M. Habegger, S. M. Hartig, A. L. Hevener, S. B. Heymsfield, C. D. Holman, M. H. de Angelis, D. E. James, L. Kazak, J. B. Kim, M. Klingenspor, X. Kong, S. Kooijman, L. Lantier, K. C. K. Lloyd, J. C. Lo, I. J. Lodhi, P. S. MacLean, O. P. McGuinness, G. Medina-Gómez, R. G. Mirmira, C. D. Morrison, G. J. Morton, T. D. Müller, Y. Ogawa, D. Pajuelo-Reguera, M. J. Potthoff, N. Qi, M. L. Reitman, P. C. N. Rensen, J. Rozman, J. M. Rutkowsky, K. Sakamoto, P. E. Scherer, G. J. Schwartz, R. Sedlacek, M. Selloum, S. R. Shaikh, S. Chen, G. I. Shulman, V. Škop, A. A. Soukas, J. R. Speakman, B. M. Spiegelman, G. R. Steinberg, K. J. Svensson, J. P. Thyfault, T. Tiganis, P. M. Titchenell, N. Turner, L. A. Velloso, A. Vidal-Puig, C. S. Ward, A. S. Williams, C. Wolfrum, A. W. Xu, Y. Xu, J. R. Zierath, International Indirect Calorimetry Consensus Committee (IICCC), A consensus guide to preclinical indirect calorimetry experiments. Nat. Metab. 7, 1765–1780 (2025).

49. E. Fabbrini, B. S. Mohammed, F. Magkos, K. M. Korenblat, B. W. Patterson, S. Klein, Alterations in adipose tissue and hepatic lipid kinetics in obese men and women with nonalcoholic fatty liver disease. Gastroenterology 134, 424–431 (2008).

50. S. B. Heymsfield, L. J. Aronne, P. Montgomery, L. B. Klickstein, L. A. Coleman, K. Dole, L. Mindeholm, S. Spruill, X. Li, K. M. Attie, BELIEVE trial investigators, Bimagrumab plus semaglutide alone or in combination for the treatment of obesity: a randomized phase 2 trial. Nat. Med. 32, 869–882 (2026).

51. E. Nunn, N. Jaiswal, M. Gavin, K. Uehara, M. Stefkovich, K. Drareni, R. Calhoun, M. Lee, C. D. Holman, J. A. Baur, P. Seale, P. M. Titchenell, Antibody blockade of activin type II receptors preserves skeletal muscle mass and enhances fat loss during GLP-1 receptor agonism. Mol. Metab. 80, 101880 (2024).

52. M. Carlsson, E. Frank, J. M. Màrmol, M. S. Ali, S. H. Raun, E. Battey, N. R. Andersen, A. Irazoki, C. Lund, C. Henríquez-Olguin, M. K. Højfeldt, P. Blomquist, F. D. Bromer, G. Mocciaro, A. Lodberg, C. B. Folsted Andersen, M. Eijken, A. M. Fritzen, J. R. Knudsen, E. A. Richter, L. Sylow, Activin receptor type IIA/IIB blockade increases muscle mass and strength, but compromises glycemic control in mice. Mol. Metab. 102, 102261 (2025).

53. M. Vacca, I. Kamzolas, L. M. Harder, F. Oakley, C. Trautwein, M. Hatting, T. Ross, B. Bernardo, A. Oldenburger, S. T. Hjuler, I. Ksiazek, D. Lindén, D. Schuppan, S. Rodriguez-Cuenca, M. M. Tonini, T. R. Castañeda, A. Kannt, C. M. P. Rodrigues, S. Cockell, O. Govaere, A. K. Daly, M. Allison, K. Honnens de Lichtenberg, Y. O. Kim, A. Lindblom, S. Oldham, A.-C. Andréasson, F. Schlerman, J. Marioneaux, A. Sanyal, M. B. Afonso, R. Younes, Y. Amano, S. L. Friedman, S. Wang, D. Bhattacharya, E. Simon, V. Paradis, A. Burt, I. M. Grypari, S. Davies, A. Driessen, H. Yashiro, S. Pors, M. Worm Andersen, M. Feigh, C. Yunis, P. Bedossa, M. Stewart, H. L. Cater, S. Wells, J. M. Schattenberg, Q. M. Anstee, Q. M. Anstee, A. K. Daly, S. Cockell, D. Tiniakos, P. Bedossa, A. Burt, F. Oakley, H. J. Cordell, C. P. Day, K. Wonders, P. Missier, M. McTeer, L. Vale, Y. Oluboyede, M. Breckons, J. Boyle, P. M. Bossuyt, H. Zafarmand, Y. Vali, J. Lee, M. Nieuwdorp, A. G. Holleboom, A. Angelakis, J. Verheij, V. Ratziu, K. Clément, R. Patino-Navarrete, R. Pais, V. Paradis, D. Schuppan, J. M. Schattenberg, R. Surabattula, S. Myneni, Y. O. Kim, B. K. Straub, A. Vidal-Puig, M. Vacca, S. Rodrigues-Cuenca, M. Allison, I. Kamzolas, E. Petsalaki, M. Campbell, C. J. Lelliott, S. Davies, M. Orešič, T. Hyötyläinen, A. McGlinchey, J. M. Mato, Ó. Millet, J.-F. Dufour, A. Berzigotti, M. Masoodi, N. F. Lange, M. Pavlides, S. Harrison, S. Neubauer, J. Cobbold, F. Mozes, S. Akhtar, S. Olodo-Atitebi, R. Banerjee, E. Shumbayawonda, A. Dennis, A. Andersson, I. Wigley, M. Romero-Gómez, E. Gómez-González, J. Ampuero, J. Castell, R. Gallego-Durán, I. Fernández-Lizaranzu, R. Montero-Vallejo, M. Karsdal, D. G. K. Rasmussen, D. J. Leeming, A. Sinisi, K. Musa, E. Sandt, M. M. Tonini, E. Bugianesi, C. Rosso, A. Armandi, F. Marra, A. Gastaldelli, G. Svegliati, J. Boursier, S. Francque, L. Vonghia, A. Verrijken, E. Dirinck, A. Driessen, M. Ekstedt, S. Kechagias, H. Yki-Järvinen, K. Porthan, J. Arola, S. van Mil, G. Papatheodoridis, H. Cortez-Pinto, A. P. Silva, C. M. P. Rodrigues, L. Valenti, S. Pelusi, S. Petta, G. Pennisi, L. Miele, A. Liguori, A. Geier, M. Rau, C. Trautwein, J. Reißing, G. P. Aithal, S. Francis, N. Palaniyappan, C. Bradley, P. Hockings, M. Schneider, P. N. Newsome, S. Hübscher, D. Wenn, J. Magnanensi, A. Trylesinski, R. Mayo, C. Alonso, K. Duffin, J. W. Perfield, Y. Chen, M. L. Hartman, C. Yunis, M. Miller, Y. Chen, E. J. McLeod, T. Ross, B. Bernardo, C. Schölch, J. Ertle, R. Younes, H. Coxson, E. Simon, J. Gogain, R. Ostroff, L. Alexander, H. Biegel, M. S. Kjær, L. M. Harder, N. Al-Sari, S. S. Veidal, A. Oldenburger, J. Ellegaard, M.-M. Balp, L. Jennings, M. Martic, J. Löffler, D. Applegate, R. Torstenson, D. Lindén, C. Fournier-Poizat, A. Llorca, M. Kalutkiewicz, K. Pepin, R. Ehman, G. Horan, G. Ho, D. Tai, E. Chng, T. Xiao, S. D. Patterson, A. Billin, L. Doward, J. Twiss, P. Thakker, Z. Derdak, H. Yashiro, H. Landgren, C. Lackner, A. Gouw, P. Hytiroglou, O. Govaere, C. Brass, D. Tiniakos, J. W. Perfield, E. Petsalaki, P. Davidsen, A. Vidal-Puig, L. I. The, An unbiased ranking of murine dietary models based on their proximity to human metabolic dysfunction-associated steatotic liver disease (MASLD). Nat. Metab. 6, 1178– 1196 (2024).

54. J. W. Mastaitis, D. Gomez, J. G. Raya, D. Li, S. Min, M. Stec, S. Kleiner, T. McWilliams, J. Y. Altarejos, A. J. Murphy, G. D. Yancopoulos, M. W. Sleeman, GDF8 and activin A blockade protects against GLP-1–induced muscle loss while enhancing fat loss in obese male mice and non-human primates. Nat. Commun. 16, 4377 (2025).

55. E. Latres, J. Mastaitis, W. Fury, L. Miloscio, J. Trejos, J. Pangilinan, H. Okamoto, K. Cavino, E. Na, A. Papatheodorou, Activin A more prominently regulates muscle mass in primates than does GDF8. Nat. Commun. 8, 15153 (2017).

56. T. Puolakkainen, P. Rummukainen, V. Pihala-Nieminen, O. Ritvos, E. Savontaus, R. Kiviranta, Treatment with Soluble Activin Type IIB Receptor Ameliorates Ovariectomy-Induced Bone Loss and Fat Gain in Mice. Calcif. Tissue Int. 110, 504–517 (2022).

57. Q. Wang, T. Guo, J. Portas, A. C. McPherron, A soluble activin receptor type IIB does not improve blood glucose in streptozotocin-treated mice. Int J Biol Sci 11, 199–208 (2015).

58. M. Thomas, B. Langley, C. Berry, M. Sharma, S. Kirk, J. Bass, R. Kambadur, Myostatin, a negative regulator of muscle growth, functions by inhibiting myoblast proliferation. J Biol Chem 275, 40235–43 (2000).

59. A. J. Levine, A. H. Brivanlou, GDF3, a BMP inhibitor, regulates cell fate in stem cells and early embryos. Development 133, 209–216 (2006).

60. M. Shiozaki, R. Sakai, M. Tabuchi, T. Nakamura, K. Sugino, H. Sugino, Y. Eto, Evidence for the participation of endogenous activin A/erythroid differentiation factor in the regulation of erythropoiesis. Proc. Natl. Acad. Sci. U. S. A. 89, 1553–1556 (1992).

61. J. J. Shen, L. Huang, L. Li, C. Jorgez, M. M. Matzuk, C. W. Brown, Deficiency of growth differentiation factor 3 protects against diet-induced obesity by selectively acting on white adipose. Mol. Endocrinol. Baltim. Md 23, 113–123 (2009).

62. I. H. Jang, A. Carey, V. Kruglov, K. Nguyen, J. R. Misialek, S. H. Cholensky, D. M. Smith, S. Bai, T. Nottoli, D. A. Bernlohr, P. L. Lutsey, C. D. Camell, GDF3 promotes adipose tissue macrophage-mediated inflammation via altered chromatin accessibility during aging. *Nat*. Aging 6, 127–142 (2026).

63. P. Arner, Catecholamine-induced lipolysis in obesity. Int. J. Obes. Relat. Metab. Disord. 23 (1999).

64. X. Zhang, S. S. Panicker, J. M. Bollinger, A. Majumdar, R. Kheireddine, L. F. Dabill, C. Kim, B. Kleiboeker, F. Zhang, Y. Chen, K. L. Magee, B. S. Learman, A. Kepecs, G. A. Meyer, J. Liu, S. A. Thomas, I. J. Lodhi, O. A. MacDougald, E. L. Scheller, A catecholamine-independent pathway controlling adaptive adipocyte lipolysis. Nat. Metab. 8, 96–115 (2026).

65. H. Wahrenberg, F. Lönnqvist, P. Arner, Mechanisms underlying regional differences in lipolysis in human adipose tissue. J. Clin. Invest. 84, 458 (1989).

66. A. J. Levine, A. H. Brivanlou, GDF3 at the crossroads of TGF-beta signaling. Cell Cycle Georget. Tex 5, 1069–1073 (2006).

67. Q. Li, X. Liu, Y. Wu, J. An, S. Hexige, Y. Ling, M. Zhang, X. Yang, L. Yu, The conditioned medium from a stable human GDF3-expressing CHO cell line, induces the differentiation of PC12 cells. Mol. Cell. Biochem. 359, 115–123 (2012).

68. P. J. Ferrara, M. J. Lang, J. M. Johnson, S. Watanabe, K. L. McLaughlin, J. A. Maschek, A. R. P. Verkerke, P. Siripoksup, A. Chaix, J. E. Cox, K. H. Fisher-Wellman, K. Funai, Weight loss increases skeletal muscle mitochondrial energy efficiency in obese mice. Life Metab. 2, load014 (2023).

69. I. Hirayama, Z. Yi, S. Izumi, I. Arai, W. Suzuki, Y. Nagamachi, H. Kuwano, T. Takeuchi, T. Izumi, Genetic analysis of obese diabetes in the TSOD mouse. Diabetes 48, 1183–1191 (1999).

70. C. F. Ibáñez, Comment on Bu et al. Insulin Regulates Lipolysis and Fat Mass by Upregulating Growth/Differentiation Factor 3 in Adipose Tissue Macrophages. Diabetes 2018;67:1761–1772. *Diabetes* **67**, e1 (2018).

71. T. Izumi, Response to Comment on Bu et al. Insulin Regulates Lipolysis and Fat Mass by Upregulating Growth/Differentiation Factor 3 in Adipose Tissue Macrophages. Diabetes 2018;67:1761–1772. *Diabetes* **67**, e2–e3 (2018).

72. A. C. McPherron, S.-J. Lee, Suppression of body fat accumulation in myostatin-deficient mice. J Clin Invest 109, 595–601 (2002).

73. J. Lin, H. B. Arnold, M. A. Della-Fera, M. J. Azain, D. L. Hartzell, C. A. Baile, Myostatin Knockout in Mice Increases Myogenesis and Decreases Adipogenesis. Biochem. Biophys. Res. Commun. 291, 701–706 (2002).

74. K. Sjöholm, J. Palming, T. C. Lystig, E. Jennische, T. K. Woodruff, B. Carlsson, L. M. S. Carlsson, The expression of inhibin beta B is high in human adipocytes, reduced by weight loss, and correlates to factors implicated in metabolic disease. Biochem. Biophys. Res. Commun. 344, 1308–1314 (2006).

75. B. J. Feldman, R. S. Streeper, R. V. Farese, K. R. Yamamoto, Myostatin modulates adipogenesis to generate adipocytes with favorable metabolic effects. Proc. Natl. Acad. Sci. U. S. A. 103, 15675–15680 (2006).

76. P. Bertolino, R. Holmberg, E. Reissmann, O. Andersson, P.-O. Berggren, C. F. Ibáñez, Activin B receptor ALK7 is a negative regulator of pancreatic beta-cell function. Proc. Natl. Acad. Sci. U. S. A. 105, 7246–7251 (2008).

77. B. Magnusson, P.-A. Svensson, L. M. S. Carlsson, K. Sjöholm, Activin B inhibits lipolysis in 3T3-L1 adipocytes. Biochem. Biophys. Res. Commun. 395, 373–376 (2010).

78. M. L. Brown, A. Lopez, N. Meyer, A. Richter, T. B. Thompson, FSTL3-Neutralizing Antibodies Enhance Glucose-Responsive Insulin Secretion in Dysfunctional Male Mouse and Human Islets. Endocrinology 162, bqab019 (2021).

79. N. Kobayashi, Y. Okazaki, A. Iwane, K. Hara, M. Horikoshi, M. Awazawa, K. Soeda, M. Matsushita, T. Sasako, K. Yoshimura, N. Itoh, K. Kobayashi, Y. Seto, T. Yamauchi, H. Aburatani, M. Blüher, T. Kadowaki, K. Ueki, Activin B improves glucose metabolism via induction of Fgf21 and hepatic glucagon resistance. Nat. Commun. 16, 3678 (2025).

80. I. Akpan, M. D. Goncalves, R. Dhir, X. Yin, E. Pistilli, S. Bogdanovich, T. Khurana, J. Ucran, J. Lachey, R. S. Ahima, The effects of a soluble activin type IIB receptor on obesity and insulin sensitivity. Int. J. Obes. 2005 33, 1265–1273 (2009).

81. S. Aykul, J. Maust, V. Thamilselvan, M. Floer, E. Martinez-Hackert, Smad2/3 Activation Regulates Smad1/5/8 Signaling via a Negative Feedback Loop to Inhibit 3T3-L1 Adipogenesis. Int. J. Mol. Sci. 22, 8472 (2021).

82. R. Kumar, A. V. Grinberg, H. Li, T.-H. Kuo, D. Sako, L. Krishnan, K. Liharska, J. Li, R. Grenha, M. C. Maguire, S. D. Briscoe, R. S. Pearsall, B. R. Herrin, R. N. V. S. Suragani, R. Castonguay, Functionally diverse heteromeric traps for ligands of the transforming growth factor-β superfamily. Sci. Rep. 11, 18341 (2021).

83. E. Fryk, J. Olausson, K. Mossberg, L. Strindberg, M. Schmelz, H. Brogren, L.-M. Gan, S. Piazza, A. Provenzani, B. Becattini, L. Lind, G. Solinas, P.-A. Jansson, Hyperinsulinemia and insulin resistance in the obese may develop as part of a homeostatic response to elevated free fatty acids: A mechanistic case-control and a population-based cohort study. EBioMedicine 65, 103264 (2021).

84. P. Petrus, N. Mejhert, P. Corrales, S. Lecoutre, Q. Li, E. Maldonado, A. Kulyte, Y. Lopez, M. Campbell, J. R. Acosta, Transforming growth factor-β3 regulates adipocyte number in subcutaneous white adipose tissue. Cell Rep 25, 551–560. e5 (2018).

85. F. Boel, V. Akimov, M. Teuchler, M. K. Terkelsen, C. W. Wernberg, F. T. Larsen, P. Hallenborg, M. M. Lauridsen, A. Krag, S. Mandrup, Deep proteome profiling of metabolic dysfunction-associated steatotic liver disease. Commun. Med. 5, 56 (2025).

86. F. Darbellay, A. Necsulea, Comparative Transcriptomics Analyses across Species, Organs, and Developmental Stages Reveal Functionally Constrained lncRNAs. Mol. Biol. Evol. 37, 240–259 (2020).

87. L. A. Broadfield, J. A. G. Duarte, R. Schmieder, D. Broekaert, K. Veys, M. Planque, K. Vriens, Y. Karasawa, F. Napolitano, S. Fujita, M. Fujii, M. Eto, B. Holvoet, R. Vangoitsenhoven, J. Fernandez-Garcia, J. Van Elsen, J. Dehairs, J. Zeng, J. Dooley, R. A. Rubio, J. van Pelt, T. G. P. Grünewald, A. Liston, C. Mathieu, C. M. Deroose, J. V. Swinnen, D. Lambrechts, D. di Bernardo, S. Kuroda, K. De Bock, S.-M. Fendt, Fat Induces Glucose Metabolism in Nontransformed Liver Cells and Promotes Liver Tumorigenesis. Cancer Res. 81, 1988–2001 (2021).

88. A. Loft, A. J. Alfaro, S. F. Schmidt, F. B. Pedersen, M. K. Terkelsen, M. Puglia, K. K. Chow, A. Feuchtinger, M. Troullinaki, A. Maida, G. Wolff, M. Sakurai, R. Berutti, B. Ekim Üstünel, P. Nawroth, K. Ravnskjaer, M. B. Diaz, B. Blagoev, S. Herzig, Liver-fibrosis-activated transcriptional networks govern hepatocyte reprogramming and intra-hepatic communication. Cell Metab. 33, 1685–1700.e9 (2021).

89. S. H. Park, S.-M. Lee, Y.-J. Kim, S. Kim, ChARM: Discovery of combinatorial chromatin modification patterns in hepatitis B virus X-transformed mouse liver cancer using association rule mining. BMC Bioinformatics 17, 452 (2016).

90. C. H. Holland, R. O. Ramirez Flores, M. Myllys, R. Hassan, K. Edlund, U. Hofmann, R. Marchan, C. Cadenas, J. Reinders, S. Hoehme, A.-L. Seddek, S. Dooley, V. Keitel, P. Godoy, B. Begher-Tibbe, C. Trautwein, C. Rupp, S. Mueller, T. Longerich, J. G. Hengstler, J. Saez-Rodriguez, A. Ghallab, Transcriptomic Cross-Species Analysis of Chronic Liver Disease Reveals Consistent Regulation Between Humans and Mice. Hepatol. Commun. 6, 161–177 (2022).

91. P. Molina-Sánchez, M. Ruiz de Galarreta, M. A. Yao, K. E. Lindblad, E. Bresnahan, E. Bitterman, T. C. Martin, T. Rubenstein, K. Nie, J. Golas, S. Choudhary, M. Bárcena-Varela, A. Elmas, V. Miguela, Y. Ding, Z. Kan, L. T. Grinspan, K.-L. Huang, R. E. Parsons, D. J. Shields, R. A. Rollins, A. Lujambio, Cooperation Between Distinct Cancer Driver Genes Underlies Intertumor Heterogeneity in Hepatocellular Carcinoma. Gastroenterology 159, 2203–2220.e14 (2020).

92. S. Shalapour, X.-J. Lin, I. N. Bastian, J. Brain, A. D. Burt, A. A. Aksenov, A. F. Vrbanac, W. Li, A. Perkins, T. Matsutani, Z. Zhong, D. Dhar, J. A. Navas-Molina, J. Xu, R. Loomba, M. Downes, R. T. Yu, R. M. Evans, P. C. Dorrestein, R. Knight, C. Benner, Q. M. Anstee, M. Karin, Inflammation-induced IgA+ cells dismantle anti-liver cancer immunity. Nature 551, 340–345 (2017).

93. H. Wang, J. Lu, X. Chen, M. Schwalbe, J. E. Gorka, J. A. Mandel, J. Wang, E. S. Goetzman, S. Ranganathan, S. F. Dobrowolski, E. V. Prochownik, Acquired deficiency of peroxisomal dicarboxylic acid catabolism is a metabolic vulnerability in hepatoblastoma. J. Biol. Chem. 296, 100283 (2021).

94. A. van Koppen, L. Verschuren, A. M. van den Hoek, J. Verheij, M. C. Morrison, K. Li, H. Nagabukuro, A. Costessi, M. P. M. Caspers, T. J. van den Broek, J. Sagartz, C. Kluft, C. Beysen, C. Emson, A. J. van Gool, R. Goldschmeding, R. Stoop, I. Bobeldijk-Pastorova, S. M. Turner, G. Hanauer, R. Hanemaaijer, Uncovering a Predictive Molecular Signature for the Onset of NASH-Related Fibrosis in a Translational NASH Mouse Model. Cell. Mol. Gastroenterol. Hepatol. 5, 83–98.e10 (2018).

95. T. Tsuchida, Y. A. Lee, N. Fujiwara, M. Ybanez, B. Allen, S. Martins, M. I. Fiel, N. Goossens, H.-I. Chou, Y. Hoshida, S. L. Friedman, A simple diet-and chemical-induced murine NASH model with rapid progression of steatohepatitis, fibrosis and liver cancer. J. Hepatol. 69, 385–395 (2018).

96. M. Sultan, M. H. Schulz, H. Richard, A. Magen, A. Klingenhoff, M. Scherf, M. Seifert, T. Borodina, A. Soldatov, D. Parkhomchuk, D. Schmidt, S. O’Keeffe, S. Haas, M. Vingron, H. Lehrach, M.-L. Yaspo, A global view of gene activity and alternative splicing by deep sequencing of the human transcriptome. Science 321, 956–960 (2008).

97. J. B. Weir, New methods for calculating metabolic rate with special reference to protein metabolism. J. Physiol. 109, 1–9 (1949).

98. A. I. Mina, R. A. LeClair, K. B. LeClair, D. E. Cohen, L. Lantier, A. S. Banks, CalR: A Web-Based Analysis Tool for Indirect Calorimetry Experiments. Cell Metab 28, 656–666.e1 (2018).

99. L. Tarhan, J. Bistline, J. Chang, B. Galloway, E. Hanna, E. Weitz, Single Cell Portal: an interactive home for single-cell genomics data. BioRxiv Prepr. Serv. Biol., 2023.07.13.548886 (2023).

100. UK Biobank Whole-Genome Sequencing Consortium, Whole-genome sequencing of 490,640 UK Biobank participants. Nature 645, 692–701 (2025).

101. S. V. Eastwood, R. Mathur, M. Atkinson, S. Brophy, C. Sudlow, R. Flaig, S. de Lusignan, N. Allen, N. Chaturvedi, Algorithms for the Capture and Adjudication of Prevalent and Incident Diabetes in UK Biobank. PloS One 11, e0162388 (2016).

102. P. Cingolani, A. Platts, L. L. Wang, M. Coon, T. Nguyen, L. Wang, S. J. Land, X. Lu, D. M. Ruden, A program for annotating and predicting the effects of single nucleotide polymorphisms, SnpEff. Fly (Austin) 6, 80–92 (2012).

103. C. M. Moore, S. A. Jacobson, T. E. Fingerlin, Power and Sample Size Calculations for Genetic Association Studies in the Presence of Genetic Model Misspecification. Hum. Hered. 84, 256–271 (2019).

