## Supplemental Figures for "GDF3 is an endogenous antagonist of the ActE-ALK7/ACVR2 pathway in adipocytes"

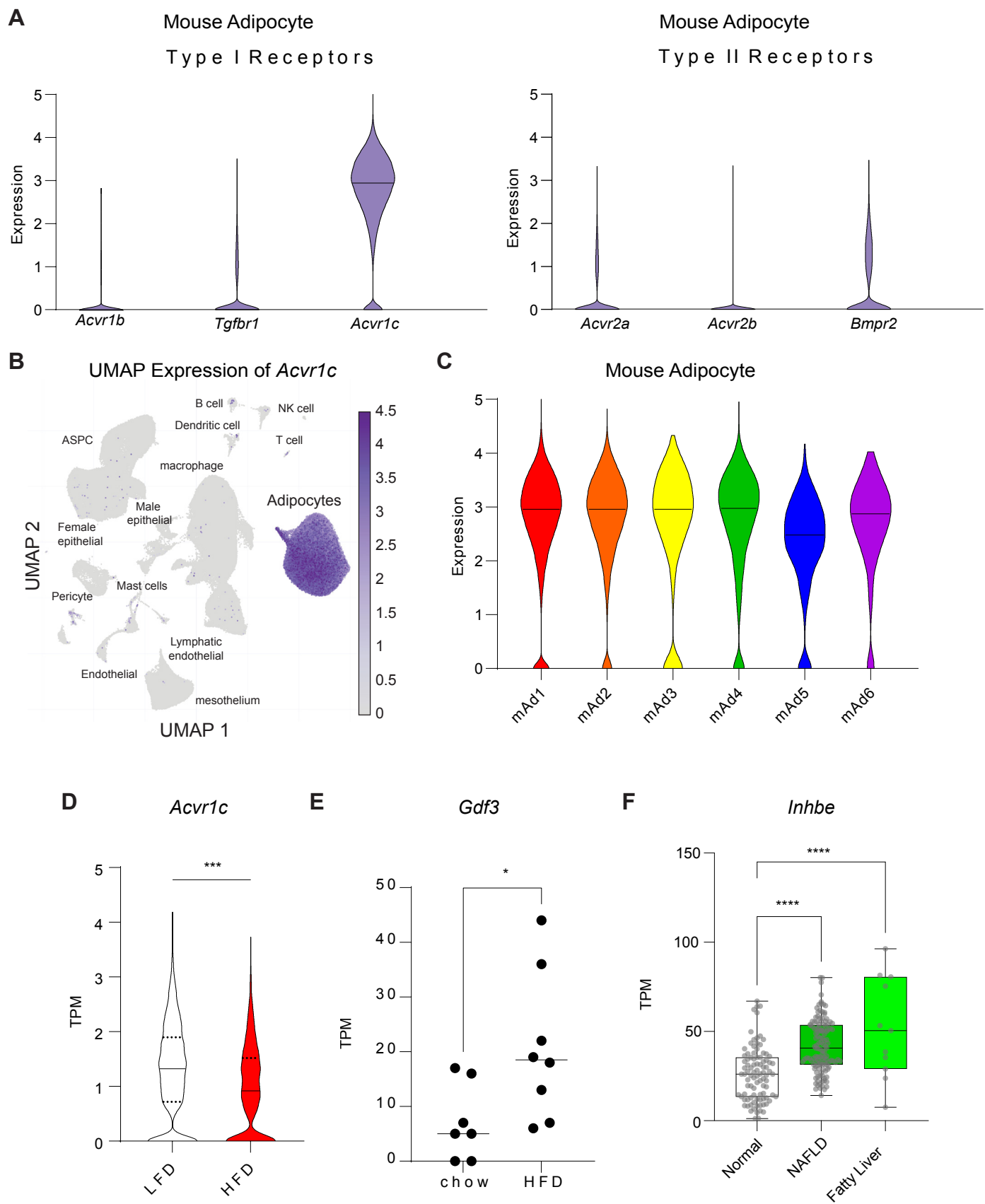

Supplementary Figure.1

### Human Adipocyte

**A**

#### Adipocyte differentiation

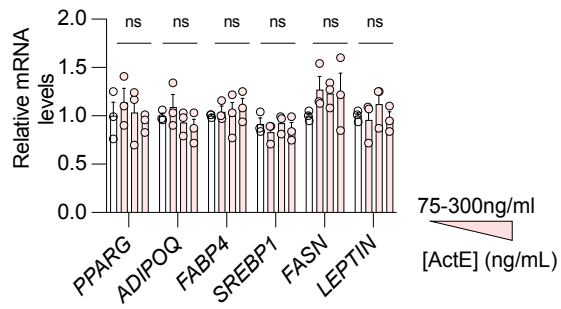

**B**

#### Adipocyte lipolysis

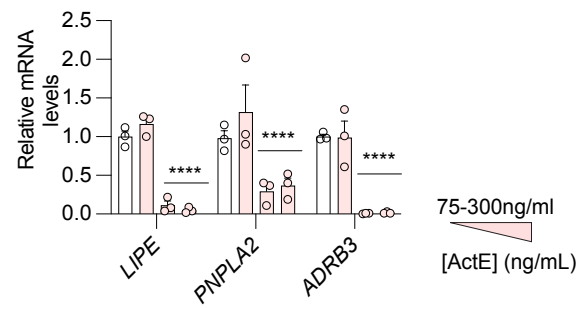

ALK5 inhibitor in mouse adipocytes

**C**

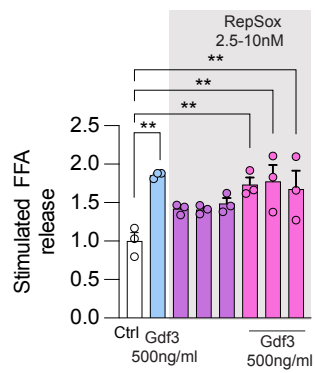

**D**

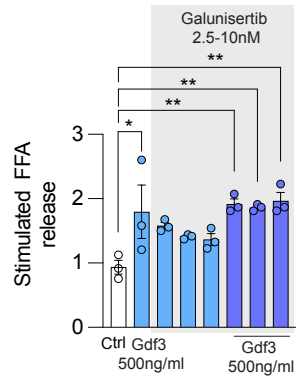

**E**

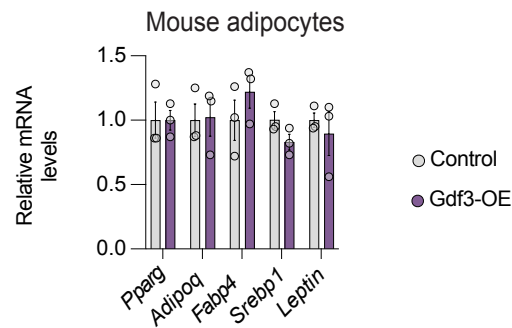

**F**

#### HEK293

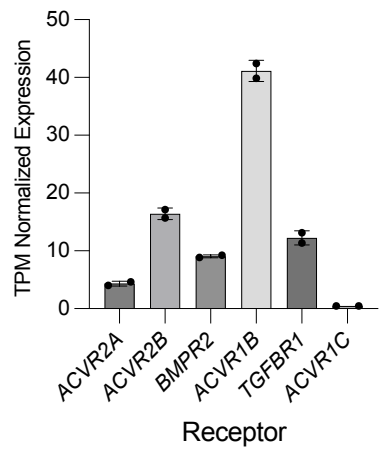

Supplementary Figure 2

**A**

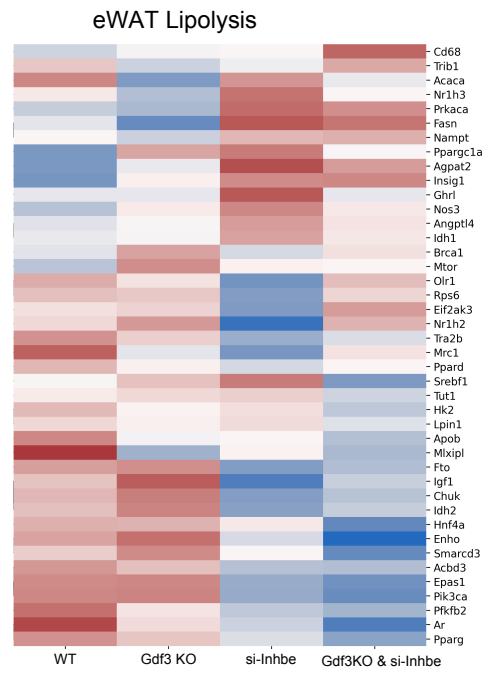

**B**

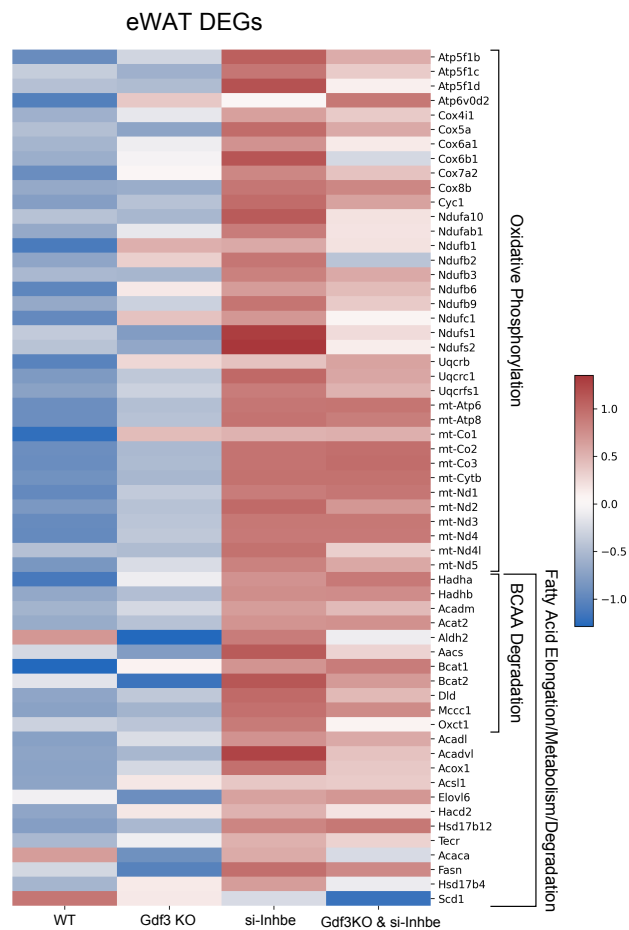

**Supplementary Figure 3**

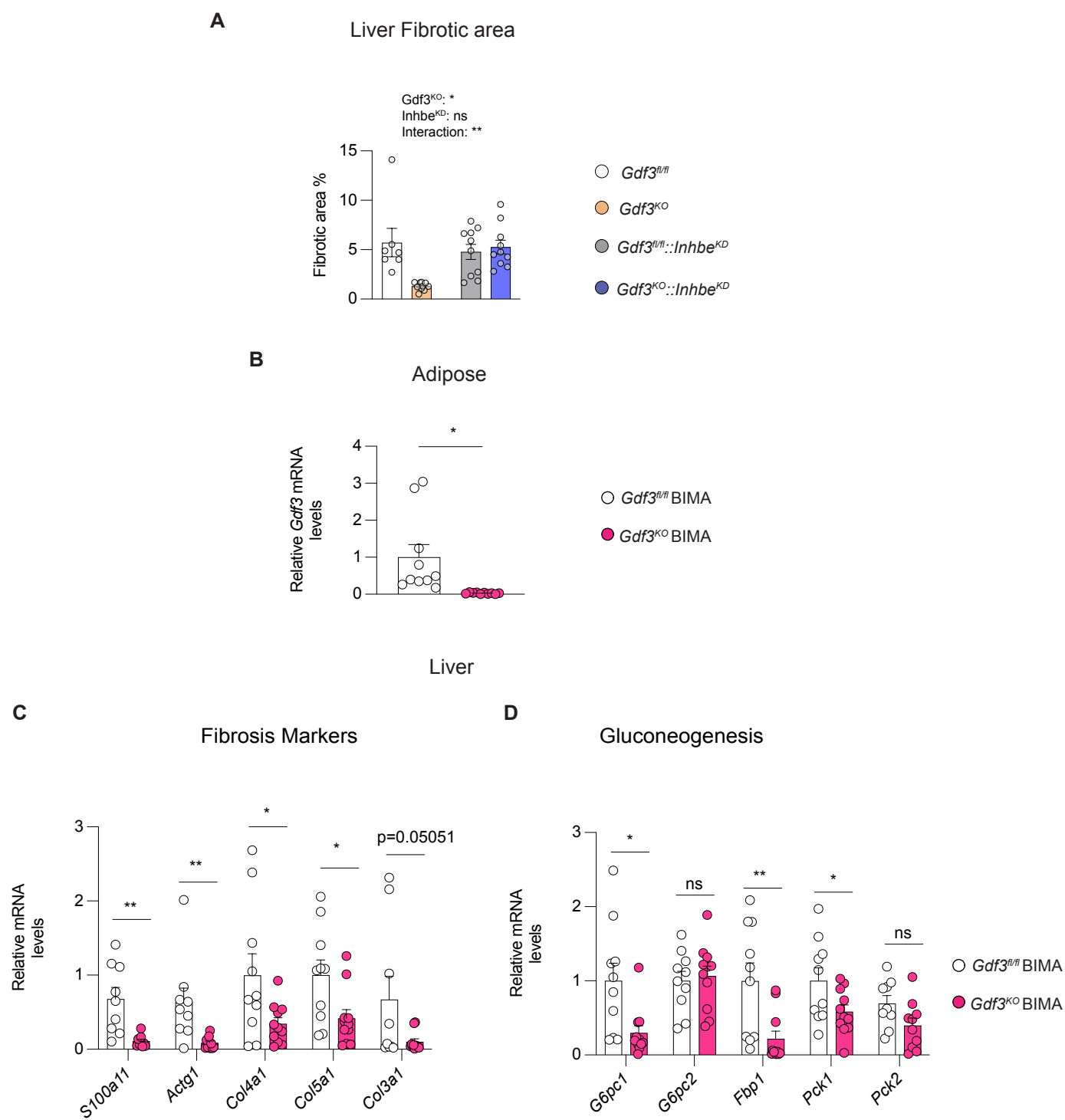

Supplementary Figure 4

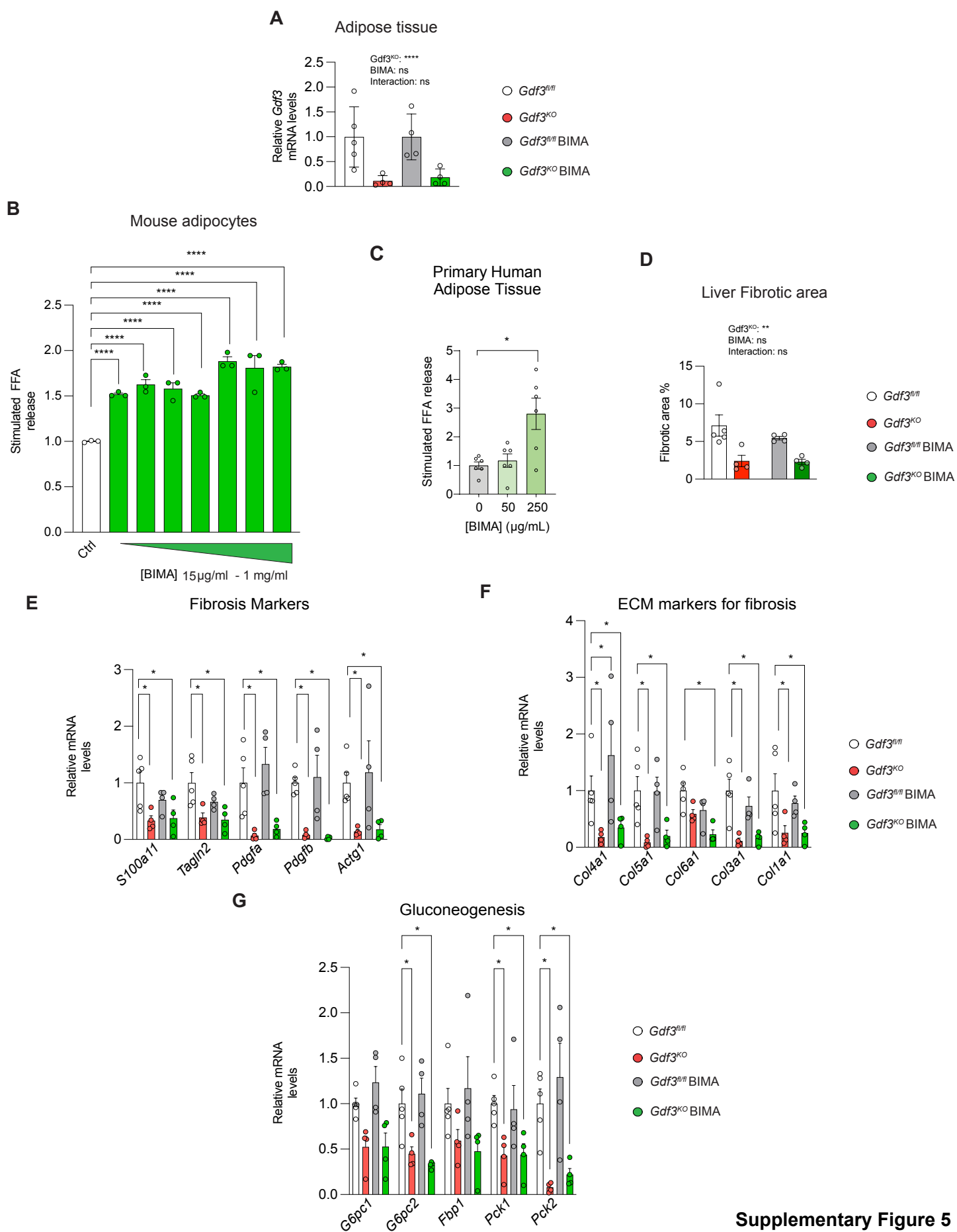

Supplementary Figure 5
